# Genomics of speciation in a great speciator (Aves: *Zosterops*) reveals the roles of both natural and sexual selection

**DOI:** 10.64898/2026.02.25.707877

**Authors:** Maëva Gabrielli, Thibault Leroy, Camille Roux, Borja Milá, Christophe Thébaud, Benoit Nabholz

## Abstract

Examining patterns of differentiation among lineage pairs at different stages of the speciation continuum can help disentangle the effects of linked and divergent selection, and identify lineage-specific targets of selection that may act as barrier loci during speciation. Here, we developed this approach using genomic data from a set of African and Indian Ocean belonging to the Zosteropidae family, commonly referred to as a “Great Speciator”. We sampled a divergence continuum aiming to identify candidate barrier loci in phenotypically, genetically distinct, and parapatrically distributed, forms from a species complex, through comparisons with closely related species. Using three independent approaches, we showed that divergence at selected loci between geographic forms is mainly concentrated on the Z chromosome, unless forms differ in their ecologies. Functional annotation revealed that genes linked to song formation and immunity could be involved in reproductive isolation between forms only separated by physical barriers, while it is mostly genes related to adaptation to local conditions, morphology, and song behaviour that may explain reproductive isolation between forms with different ecologies, highlighting a role for combined effects of natural and sexual selection in driving the evolution of reproductive barriers in this rapidly speciating lineage

## Introduction

Identifying candidate genomic regions that may contribute to barriers to gene flow between diverging populations is a key step in speciation genomic studies (Feder et al. 2012; Ravinet et al. 2017). To that aim, the analysis of genomic landscapes of differentiation, typically measured by the fixation index *F_ST_* (Wright 1943), has been widely used in a wide array of organisms, such as plants (Renaut et al. 2013; Stankowski et al. 2019; Leroy et al. 2020; Shang et al. 2023), insects (Turner et al. 2005; Nadeau et al. 2012; Soria-Carrasco et al. 2014), fishes (Marques et al. 2016; Duranton et al. 2018), mammals (Harr 2006; Carneiro et al. 2010) and birds (Ellegren et al. 2012; Poelstra et al. 2014; Burri et al. 2015; Vijay et al. 2016; Irwin et al. 2018; Sendell-Price et al. 2020). A consistent pattern that emerged from these studies is the very large heterogeneity in the levels of differentiation along genomes, where regions of low differentiation coexist alongside regions that are broadly differentiated (Cruickshank and Hahn 2014; Ravinet et al. 2017). In addition, the biological interpretation of highly differentiated regions was found to be highly dependent on a number of underlying assumptions (e.g., demographic history and genomic architecture of divergence; Burri 2017; Ellegren and Wolf 2017; Ravinet et al. 2017; Rodrigues et al. 2024; Schield et al. 2025), challenging the view that peaks of differentiation can be readily interpreted as ‘islands of speciation’ (Turner et al. 2005), in which a local reduction of gene flow is due to divergent selection (Feder et al. 2012). Several studies have also shown that heterogeneous differentiation across the genome can reflect the effects of background selection (Charlesworth et al. 1993; Noor and Bennett 2009; Turner and Hahn 2010; Cruickshank and Hahn 2014; Booker et al. 2022; Liang et al. 2022) as well as endogenous selection on intrinsic genetic incompatibilities (Bierne et al. 2011; Ravinet et al. 2017). In addition, the effects of different adaptive and non-adaptive processes on *F_ST_* extend over wider genomic regions when local recombination is reduced (e.g., Vijay et al. 2016), potentially impacting the neighbouring neutral regions on large chromosomal portions (Burri et al. 2015). This important effect of local variation in recombination rates can vary across species and lineages, ultimately depending on how recombination landscapes evolve (Talbi et al. 2025). It is therefore essential to interpret genomic landscapes of differentiation in the light of their landscapes of recombination in order to identify genuine signals of barrier loci (Ravinet et al. 2017) and understand population divergence and speciation better.

When examining genomic differentiation landscapes, one general way to separate the effects of background selection from those due to barrier loci is to use a comparative genomic approach using a set of lineages, such as relatively closely related species, with highly conserved recombination landscapes (Kawakami et al. 2014, 2017; Singhal et al. 2015). Contrasting patterns of differentiation among populations and species pairs at different stages of the speciation continuum but sharing a recombination landscape can help disentangle the effects of linked and divergent selection, and identify lineage-specific targets of selection that may act as barrier loci (Roux et al. 2016; Burri 2017). In this context, the choice of the appropriate evolutionary controls is crucial, with respect to both the stage of differentiation within each pair of populations or species and the phylogenetic divergence between pairwise comparisons (Burri 2017). Comparing parapatric and allopatric population pairs, where gene flow is possible or limited (see Marques et al. 2016) in systems with a well-annotated reference genome and a well-characterized evolutionary history may provide a robust framework to identify speciation gene candidates.

Here, we analyze genomic landscapes of differentiation to identify and characterize candidate barrier loci that may contribute to reproductive isolation between geographic forms in a species complex, the Reunion Grey White-eye (*Zosterops borbonicus*). This species complex is endemic to the volcanic oceanic island of Reunion (Mascarene archipelago) and it belongs to the white-eye family, the Zosteropidae, which is represented in the southwestern Indian Ocean by several closely related species showing varying degrees of divergence relative to the Reunion Grey white-eye (Warren et al. 2006; Martins et al. 2020). It displays the typical features of a great speciator sensu Diamond et al. (1976), i.e. propensity to diversify on islands (Warren et al. 2006; Moyle et al. 2009; Milá et al. 2010; Gabrielli et al. 2020). Four genetically and phenotypically distinct forms with abutting ranges have been recognized in this complex (Gill 1973; Milá et al. 2010; Cornuault et al. 2015). They meet at narrow and stable secondary hybrid zones (Gill 1973; Bertrand et al. 2014; Delahaie et al. 2017), indicating that there must be genetic differences that prevent the populations from mixing, and consequently, some degree of reproductive isolation between the different forms (Cowles and Uy 2019). Three forms are found in the lowlands, primarily below 1,400 meters, with contact zones coinciding with natural physical barriers, such as large rivers or lava flows, that are not associated with drastic changes in environmental conditions (Delahaie et al. 2017). These lowland forms differ in plumage colour, with a brown-headed brown form (LBHB for Lowland Brown-Headed Brown) with a light brown back and head, a grey-headed brown form (GHB) with a brown back and a grey head, and a brown-naped brown form (BNB) with a brown back and nape and a grey crown (Figure 1). A fourth form (HIGH), found in the highlands between 1,400 meters and 3,000 meters, is slightly larger than the lowland forms and comprises two distinct genetically determined colour phenotypes, with birds showing predominantly grey or brown plumage, respectively (Gill 1973; Cornuault et al. 2015; Bourgeois et al. 2017; Mould et al. 2023; Figure 1). This highland form occupies a distinct elevational and ecological zone, suggesting that it may have experienced very different selection pressures from the lowland forms during its evolutionary history (Gabrielli et al. 2020). The highland form has an almost continuous distribution range but the populations occupying the two main volcanoes, i.e. the extinct Piton des Neiges volcano in the North and the active Piton de la Fournaise volcano in the South, show similar levels of differentiation to those found between lowland forms (Gabrielli et al. 2020). The Reunion Grey White-eye thus comprises geographical forms that have recently diverged (<0.5Myr) while already showing a high level of reproductive isolation, and provides a useful system with which to search for candidate genomic regions that are relevant to speciation. First, few barriers to gene flow are expected, and these barriers should be among the first genomic regions to have differentiated during the process of divergence. Second, a high quality reference genome has recently been produced (Leroy et al. 2021b), facilitating genomic analyses in this system. Third, the different forms of Reunion Grey White-eyes provide different hypotheses in terms of selection regimes involved: while natural selection may be primarily at play between lowland and highland forms (Bertrand et al. 2016), sexual selection may be primarily involved in the divergence between lowland forms (Delahaie et al. 2017). Accordingly, a recent genomic scan study based on pool-seq data has revealed that divergence peaks between lowland forms were restricted to Z chromosome while both autosomes and Z chromosome were involved for highland vs lowland comparisons (Bourgeois et al. 2020). Here, we take advantage of this natural experimental model to investigate candidate genomic regions that may contribute to reproductive isolation under different selective regimes in both autosomes and Z chromosome using whole-genome data and accounting for background selection.

**Figure 1:**
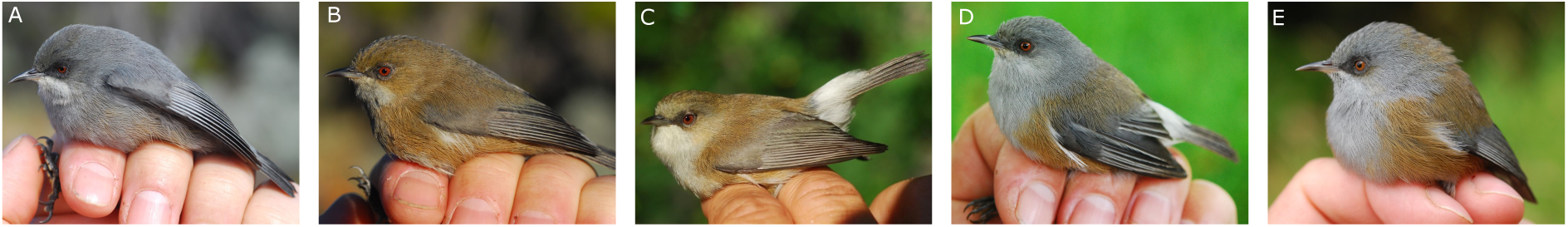
Field photographs of adult individuals representative of the different Reunion grey white-eye geographic forms: A. Grey morph from highland form (HIGH), B. Brown morph from highland form (HIGH), C. Lowland brown-headed brown form (LBHB), D. Grey-headed brown form (GHB), and E. Brown-naped brown form (BNB), Photos by Borja Milá.

Specifically, we used three approaches to identify candidate barrier loci underlying the differentiation between forms, trying to disentangle the effects of background selection from the effects of positive selection on barrier loci by: (i) identifying high differentiation outliers in each pairwise comparison, accounting for recombination rate by using the recombination landscape itself and also controlling for shared highly differentiated regions in a set of closely related species; (ii) detecting candidate loci for selective sweeps in each form; and (iii) identifying barriers to gene flow using Approximate Bayesian Computation inferences. We then combined results from these three types of analyses to characterize finely candidate genomic regions that are likely involved in the build-up of barriers to gene flow between the different forms of the Reunion Grey White-eye and to discuss how they may relate to the different selective regimes experienced by these forms.

## Methods

### a) Population sampling and DNA sequence data

We obtained blood samples from a total of 22 individuals from 22 localities (Table S1) throughout the distribution of the Reunion Grey White-eye in order to include representatives from all geographic forms. Birds were captured with mistnests, banded with uniquely numbered aluminium rings and released after processing at the capture site. To allow for evolutionary controls with respect to both differentiation stage and phylogenetic divergence, we selected other species along a speciation continuum and obtained additional blood samples from (i) Mauritius Grey White-eye (*Z. mauritianus*; nine individuals), the sister-species to the Reunion Grey White-eye (Warren et al. 2006; Milá et al. 2010), (ii) the Reunion Olive White-eye (*Z. olivaceus*; 11 individuals), sister to the clade formed by Reunion and Mauritius Grey White-eyes and endemic to Reunion (Warren et al. 2006) and (iii) the Orange River White-eye (*Z. pallidus*; two individuals) and the Cape White-eye (*Z. virens*; seven individuals), two species from mainland Africa that are more distantly related to the Reunion Grey White-eye (Cai et al. 2020; Martins et al. 2020; Leroy et al. 2021b) (Figure 2; table S1). Approximately 1 µg of high-quality DNA was extracted using a QIAGEN DNeasy Blood & Tissue kit following the manufacturer’s instructions, with an extra pre-digestion grinding step. Genomic DNA extractions were sent to Novogene Bioinformatics Technology (Beijing, China; 34 individuals) and to GeT-PlaGe core facility (INRAE, Toulouse, France; nine individuals), for shotgun whole-genome resequencing with an Illumina HiSeq2500 sequencer. Eight individuals sequenced using similar procedures were re-used from previous studies (Bourgeois et al. 2017; Leroy et al. 2021a).

**Figure 2:**
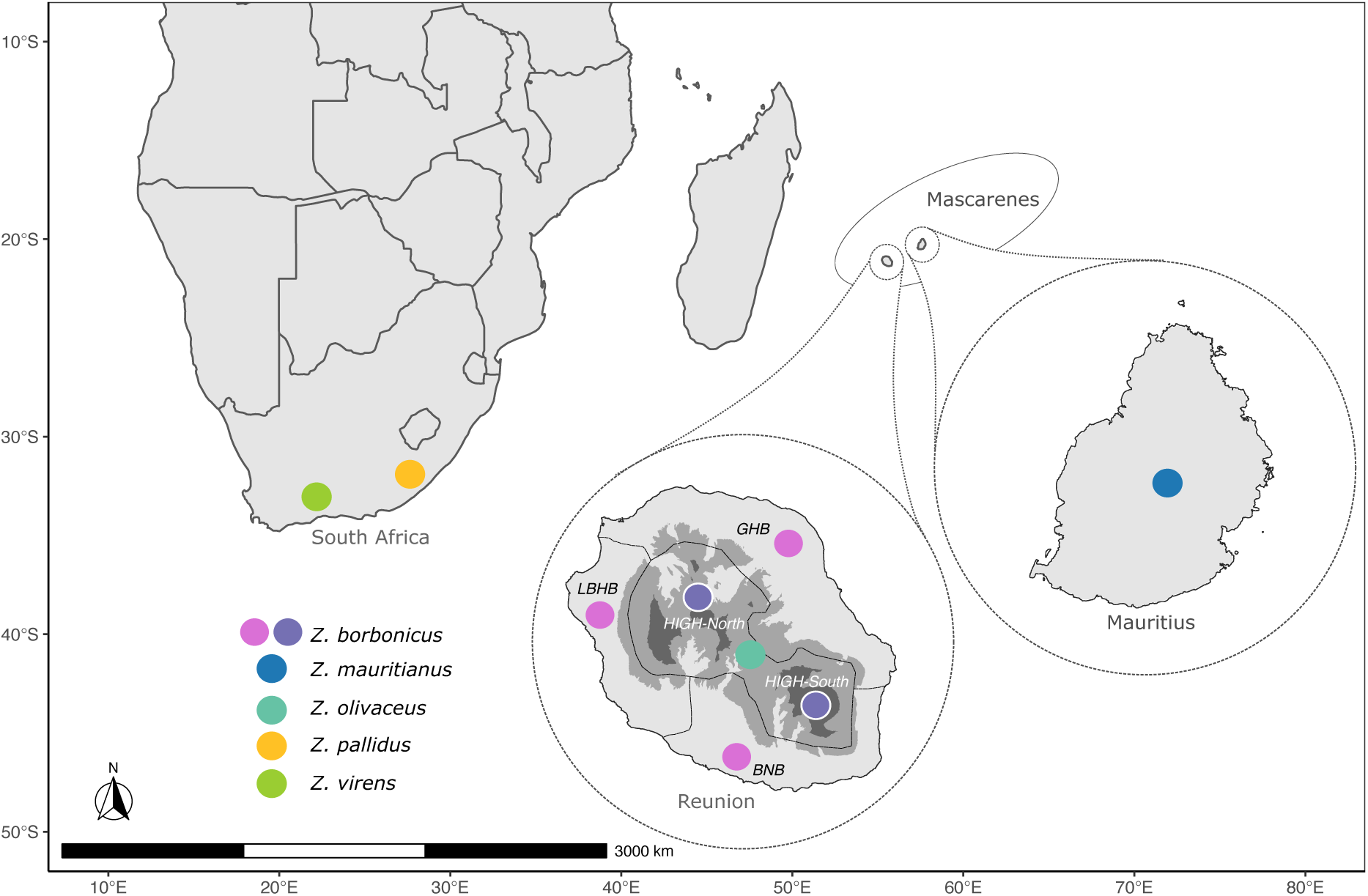
Sampling locations for the different species used in this study. In Reunion, grey colours correspond to elevation classes: light grey = elevation below 1000 m; medium grey = elevation between 1000 and 2000 m; dark grey = elevation above 2000 m.

### b) Reference-based whole-genome processing

Sequencing reads of all species were first mapped against the high-quality genome assembly (scaffold N50 exceeding one megabase) of the Reunion Grey White-eye (Leroy et al. 2021a,b) the bwa-mem algorithm (Li and Durbin 2009) with default parameters. Single nucleotide polymorphisms (SNPs) were then called for each species independently using GATK v. 3.7 (McKenna et al. 2010) using base quality score recalibration and indel realignment. Both variant and invariant sites were called (option “--includeNonVariantSites”). Variant filtration was performed using a python script to speed up computations (https://github.com/ThibaultLeroyFr/Intro2PopGenomics/blob/master/3.2.2/6-Variant_Filtration/VariantFiltrationVCF.py), but following the same procedures than under GATK, assuming the following thresholds: QD>2.0, FS<60.0, MQ>40.0, MQRANKSUM>-2.0, READPOSRANKSUM>-2.0 and RAW_MQ > 45,000. The three Mascarene species (i.e. *Z. borbonicus*, *Z. mauritianus*, and *Z. olivaceus*) were also genotyped together and using the same variant filtration procedure. We finally attributed the chromosome location for each site, based on the highly conserved synteny with Zebra Finch (*Taeniopygia guttata*) as well as with many bird species (for details, see Leroy et al. 2021a). We removed all missing data, resulting in approximately 13 million SNPs for genomic scans of differentiation between the Réunion grey white-eye and other species, and about 7 million SNPs for between-form comparisons in the Réunion grey white-eye, including selective sweep analyses.

### c) Scans of differentiation and speciation continuum

We first described the genomic landscape of differentiation by computing pairwise *F_ST_* between pairs of populations (forms) or species spanning the continuum of divergence using SNP data and windows of 50 kb using scripts from Simon Martin (https://github.com/simonhmartin/genomics_general). We then plotted distributions of *F_ST_* in autosomes and sexual chromosomes using R v. 3.2., and following the approach used by Sendell-Price et al. (2020), we calculated the skewness of *F_ST_* distribution (where μ2 and μ3 are the second and third central moments of the distribution) as a statistic to describe the *F_ST_* distribution.

### d) Correlations between *F_ST_* of Reunion Grey White-eye forms and *F_ST_* of other White-eyes from different islands and South Africa

In order to decipher the effects of background selection from positive selection, we used the observation that landscapes of differentiation can be correlated even between divergent taxa due to the effects of background selection (Doren et al. 2017; Dutoit et al. 2017; Vijay et al. 2017; Delmore et al. 2018; Irwin et al. 2018; Liang et al. 2022; Shang et al. 2023). Therefore, high differentiation between pairs of taxa in allopatry may be related to genomic architecture, while pairs of taxa in contact (parapatry) should display elevated differentiation in both genomic areas that constitute barriers to gene flow and those related to genomic architecture. Therefore, we correlated *F_ST_* between highland and lowland forms of the Reunion Grey White-eye, as well as *F_ST_* between the three Mascarene species and between both African species in order to obtain information on common regions of high *F_ST_*. Then, we plotted the recombination rate against *F_ST_* between each population (form) and species of the speciation continuum. We identified *F_ST_* outliers according to recombination rate by highlighting the top 0.5% *F_ST_* values for regions with a recombination rate higher than 1cM, for both autosomes and Z chromosome independently, and highlighted these outliers on genomic scans of differentiation as candidate loci for positive selection. Finally, we compared our candidate regions with the top 5% and 1% *F_ST_* values obtained from the other species pairs (*Z. olivaceus* - *Z. mauritianus* and *Z. virens* - *Z. pallidus*). Although these thresholds are arbitrary, they were chosen to be conservative: the most stringent threshold (0.5%) was used to identify the candidate regions, whereas the broader thresholds (1% and 5%) served as controls to assess the overlap of high-*F_ST_* regions between our focal species and the outgroups.

### e) Recombination rate analyses

We then corrected the observed local *F_ST_* values with recombination rates, as high *F_ST_* regions are often associated with low recombination due to the effects of background selection. To do so, we used the recombination landscape of the Zebra Finch obtained using multigeneration pedigrees and SNP markers (Backström et al. 2010), since recombination maps are highly conserved among birds (Kawakami et al. 2014, 2017; Singhal et al. 2015). We used loess regressions in a marey map framework (Rezvoy et al. 2007) to extrapolate local recombination rates at all positions for the Zebra Finch chromosomal assembly, for which we obtained the correspondence with our Reunion Grey White-eye assembly (see “Reference-assisted genome organization” section in Leroy et al. 2021a for details). We were able to recover local recombination rates for about 8 million SNPs.

### f) Selective sweep detection

To investigate if genomic regions of elevated differentiation have been subjected to selective sweeps, we ran Sweepfinder2 (Nielsen et al. 2005; DeGiorgio et al. 2016; Huber et al. 2016) for each geographical form of the Reunion Grey White-eye (separating the highland form into the two mountain populations HIGH-North and HIGH-South). Sweepfinder performs a genome-wide search for selective sweeps using an empirical background frequency spectrum to compute a test statistic for each site in the genome that reflects the likelihood of observing SNP data under the assumption of a recent selective sweep compared with a null model where no sweep has occurred (Composite Likelihood Ratio test, hereafter CLR).

Sweepfinder2 includes additional features to disentangle recent selective sweeps from spurious signals of background selection, including accounting for local variations in recombination rates and the *B* statistics (McVicker et al. 2009) to measure the strength of background selection. As this statistic has not been yet estimated using bird genomes, we included information about the recombination rate in the computation of the likelihood ratio.

We first ran the program using allelic frequency data only (option -s) and then added a recombination map with a pre-computed empirical spectrum based on all SNPs (including SNPs without recombination information) (option -lr). CLR was computed for 50-kb windows. We plotted the genomic scans of the CLR for each geographical form, and defined candidate selective sweeps as loci with a CLR in the top 0.1% of the CLR values.

### g) Approximate Bayesian Computation inferences of barriers to gene flow

In complement, we used an alternate strategy to identify barriers to gene flow, based on the Approximate Bayesian Computation (hereafter ABC, Csilléry et al. 2010; Pudlo et al. 2016) framework implemented in the DILS (Demographic Inferences with Linked Selection) software (Fraïsse et al. 2021). This strategy has two main advantages: (i) the inference explicitly accounts for the demography of the two populations or species that are compared, and their underlying confounding effects on genomes and (ii) the reasoning is based not only on a single summary statistic but on a combination of statistics, which is known to help differentiate patterns (e.g., Cruickshank and Hahn 2014; Shang et al. 2023). First, we inferred the timing of gene flow during divergence between two populations, using whole-genome sequencing data and accounting for potential genome-wide heterogeneity in migration and effective population size. When ongoing migration is supported to be heterogeneous, DILS can then identify regions linked to barriers to gene flow, i.e. genomic regions that have a reduced migration rate compared to the average migration rate. We ran DILS on pairs of Reunion Grey White-eye geographical forms in an attempt to identify loci that could be important for reducing gene flow between these forms. We used sequences of size ≤ 10kb with less than 50 % of missing data for all individuals to get informative loci, resulting in between 21,368 and 23,617 loci for the different pairs of the Reunion Grey White-eye geographical forms. Here, we used 10-kb loci rather than 50-kb loci to enhance our ability to detect barriers to gene flow operating in short genomic regions. Thirty-seven summary statistics were used for two population models, including estimates of genetic diversity for each population (number of biallelic sites, nucleotide diversity (π), Watterson’s *θ_W_* and Tajima’s *D*) and measures of differentiation and divergence between populations (*F_ST_*, net divergence), considering the average and standard deviation of each measure (Roux et al. 2013). For all inferences, model comparison was performed using a random forest algorithm. Two million simulations were run under the best model to perform parameter estimations, and four rounds of goodness-of-fit were run to refine parameter estimation. We used a generation time of 1 year and a mutation rate of 4.6e-9 mutations/site/generation (Smeds et al. 2016).

We used models of two-population divergence with different timing of gene flow, with two models assuming the absence of ongoing gene flow (SI: Strict Isolation; AM=Ancient Migration), and two models assuming ongoing gene flow (IM=Isolation with Migration; SC=Secondary Contact). We ran these models for three comparisons between lowland forms (GHB−LBHB; GBH−BNB; LBHB−BNB) and three comparisons between highland and lowland forms (HIGH-North−GHB; HIGH-North−LBHB; HIGH-South−BNB).

For models of heterogeneous migration along the genome, the program performs a locus-level assignment of migration rate, with a category ‘migration’ or ‘isolation’ and an associated posterior probability. We defined “barrier loci” as loci that were assigned the category ‘isolation’ with a posterior probability greater than 0.95.

### h) Gene annotations

To gain insights into the putative selective pressures acting on the different forms of the Reunion Grey White-eye, we searched for the functions of genes located in the candidate outlier regions that were identified by each of the three approaches that we used (high differentiation outliers in each pairwise comparison; candidate loci for selective sweeps in each form; and barriers to gene flow inferred with ABC for the pair HIGH-North−GHB). We retrieved gene annotations from the Zebra Finch genome (version 3.2.4; GenBank assembly accession GCA_000151805.2) within 150-kb windows centered on each candidate outlier, corresponding to one 50-kb window upstream and one downstream of the outlier window for analyses using 50-kb windows, using the intersect function in bedtools v2.25.0 (Quinlan and Hall 2010). Gene functions were manually curated based on extensive literature searches for each gene using UniProt and Google Scholar, with the queries “gene name” + adaptation and “gene name” + adaptation + bird. Annotations were assigned only when supported by experimental evidence or high-confidence functional studies, ensuring high-quality, literature-based functional classification rather than relying on automated Gene Ontology annotations.

## Results

### 1 Landscapes of differentiation across a continuum of genetic divergence

For all comparisons between populations (forms) and species, we found a heterogeneous genomic landscape of differentiation (Figure 3). Mean differentiation increased and regions of high differentiation accumulated across the genome as the extent of genetic divergence between populations or species increases (Table S2). Within Reunion Grey White-eyes, comparisons between lowland forms had lower *F_ST_* (mean autosomal *F_ST_* = 0.01) and fewer genomic peaks that comparisons between highland and lowland forms (mean autosomal *F_ST_* = 0.03), in agreement with previous studies (Delahaie et al. 2017; Bourgeois et al. 2020).

**Figure 3:**
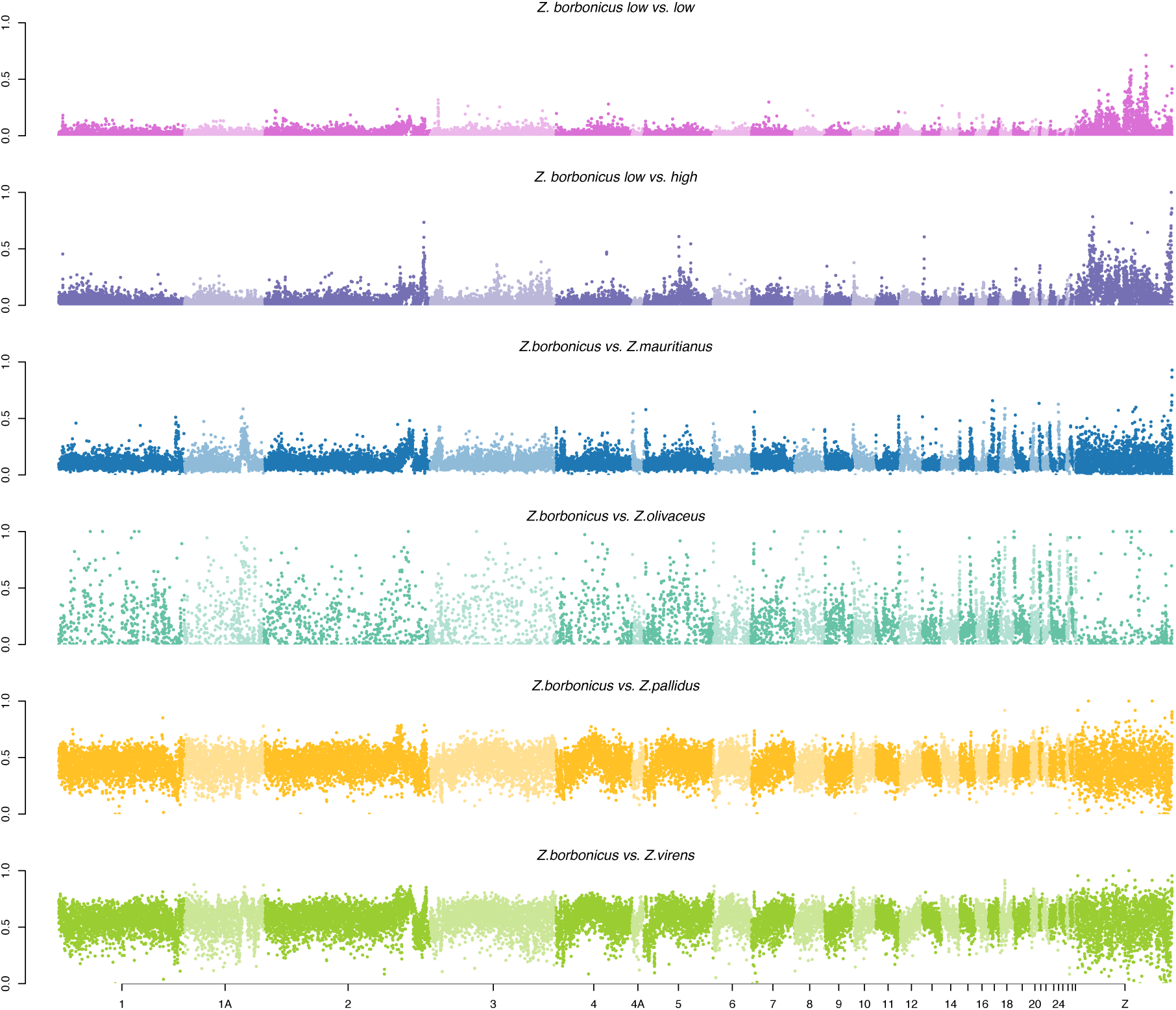
Genomic landscape of *F*_ST_ across a continuum of genetic divergence in *Zosterops*.

In comparisons between Reunion Grey White-eye geographical forms, the distribution of *F_ST_* was highly skewed towards low values of *F_ST_* (Figure S1). As expected, the skewness decreased with increasing divergence (Table S3), for both autosomes and the Z sex chromosome, reflecting the accumulation of differentiation as speciation progresses. Autosomes and the Z sex chromosome have more similar patterns of *F_ST_* distribution for higher divergence with no gene flow, and so even for pairs where the *F_ST_* is far from saturated as for comparisons with the Mauritius Grey White-eye (mean autosomal *F_ST_* = 0.08) or with the Reunion Olive White-eye (mean autosomal *F_ST_* = 0.09). Interestingly, this is not the case for cases of lower divergence with potential gene flow within Reunion Grey White-eyes, where skewness is markedly lower in sex chromosomes than in autosomes (table S3).

### 2 Candidate loci for selection in the Reunion Grey White-eye

#### 2a Searching for genomic islands of differentiation when accounting for recombination

We first explored the relationship between differentiation, measured with *F_ST_*, and recombination landscape. A negative relationship was observed between *F_ST_* and chromosome length for species comparisons (Figure S2), which can be explained by recombination as short chromosomes have higher recombination rates than large chromosomes, and *F_ST_* is negatively correlated with recombination rate (Nachman and Payseur 2012; Juric et al. 2016; Martin et al. 2019). We also tested the correlation between differentiation landscapes of Mascarene white-eyes (*Z. borbonicus*; *Z. mauritianus* and *Z. olivaceus*) and South Africa white-eyes (*Z. pallidus* and *Z. virens*). All correlations were significantly positive, confirming the combined effect of genome architecture and background selection on genomic differentiation landscape (Figure S3). This shared landscape of differentiation is also reflected in the presence of regions with elevated *F_ST_* at the end of chromosome 2 and in the central part of chromosome 4 and 5, both in the comparison between the highland and lowland forms of the Reunion Grey White-eye and in the comparison between the Reunion Grey White-eye and the South African species (Figure 3).

Not surprisingly given the correlated landscape of differentiation between distantly related species, we found a negative relationship between *F_ST_* and recombination rate estimated from the zebra finch for most comparisons and for both autosomes and sex chromosomes (Figures 4 and S4), with higher correlation coefficients for Reunion Grey White-eye comparisons (Figure 4). Strikingly, in this species, we found peaks of relatively high *F_ST_* in regions of moderate to high recombination, both in autosomes and the Z sex chromosome (Figure 4). In order to account for the possibility that linked selection could cause high differentiation in regions of low recombination, we defined genomic islands of differentiation that may have arisen though adaptive divergence as regions with very high differentiation (top 0.5% *F_ST_* values) and a moderate to high recombination rate (>1cM). This yielded up to nine genomic regions per comparison that are good candidate regions for positive selection or reduced gene flow involved in divergence (Figure 5). To confirm that these regions are not the result of the interplay between background selection and recombination rate, we verified that they fall outside the high-*F_ST_* regions (top 5% or top 1%) detected in the *Z. mauritianus* - *Z. olivaceus* and *Z. pallidus* - *Z. virens* comparisons. Only one candidate region (on chromosome 4) identified above was shared across these comparisons using the top 1% and can therefore be considered dubious. Four regions (chromosome 1A, 3, 4 and 13) were shared using the top 5%. All the candidate regions contained 87 genes based on synteny with the zebra finch (Table S4), whose functions were identified based on an extensive literature search. Overall, 0 and 24 of the identified genes fall within the high-*F_ST_* windows shared with the outgroup at the 1% and 5% thresholds, respectively (Table S4). Below, we present the biological functions of several of these genes, without emphasising any function represented solely by genes located in the shared high-*F_ST_* windows. Genomic regions of high differentiation contained loci related to physiological, immune, and behavioural processes. In lowland forms, we detected signatures of selection on genes involved in hypoxia response, energy metabolism, immune defence, pigmentation, and neurobehavioural traits, consistent with adaptation to subtle elevational, environmental, and ecological variation. Comparisons between lowland and highland forms revealed distinct signals of selection on altitude adaptation, metabolic regulation, immune function, and traits associated with communication and sensory perception, supporting adaptation to very different environmental pressures.

**Figure 4:**
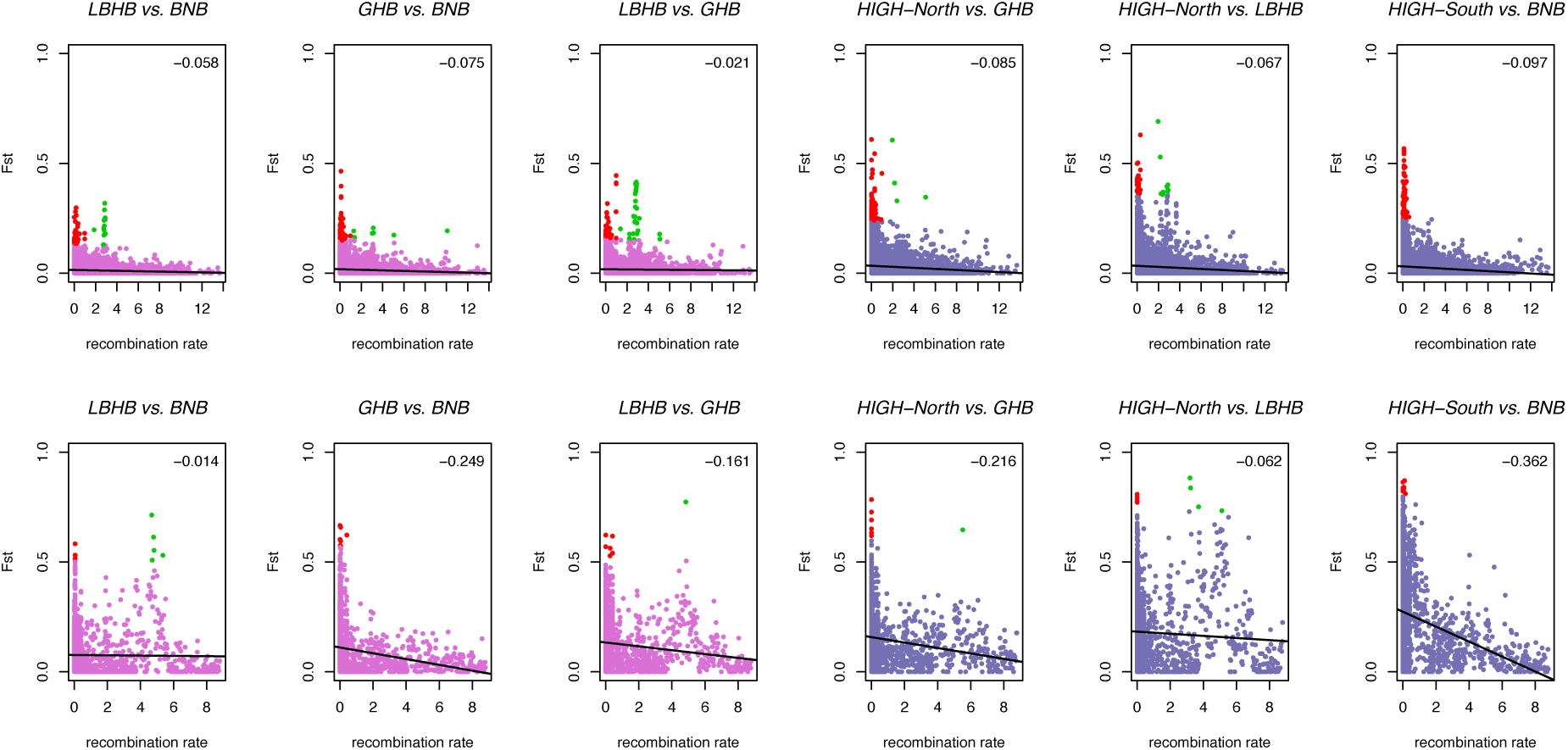
*F*_ST_ according to recombination rate in the Reunion Grey White-eye, in autosomes (top panel) and the Z chromosome (bottom panel). Top right numbers are Pearson correlation coefficients, and the black line is a linear regression. Genomic regions of high *F*_ST_ (top 0.5%) that have a low recombination rate (between 0 and 1cM), are indicated in red. Genomic regions of high *F*_ST_ (top 0.5%) with moderate to high recombination rate (>1cM), that can be viewed as candidate regions for positive selection or reduced gene flow, are indicated in green.

**Figure 5:**
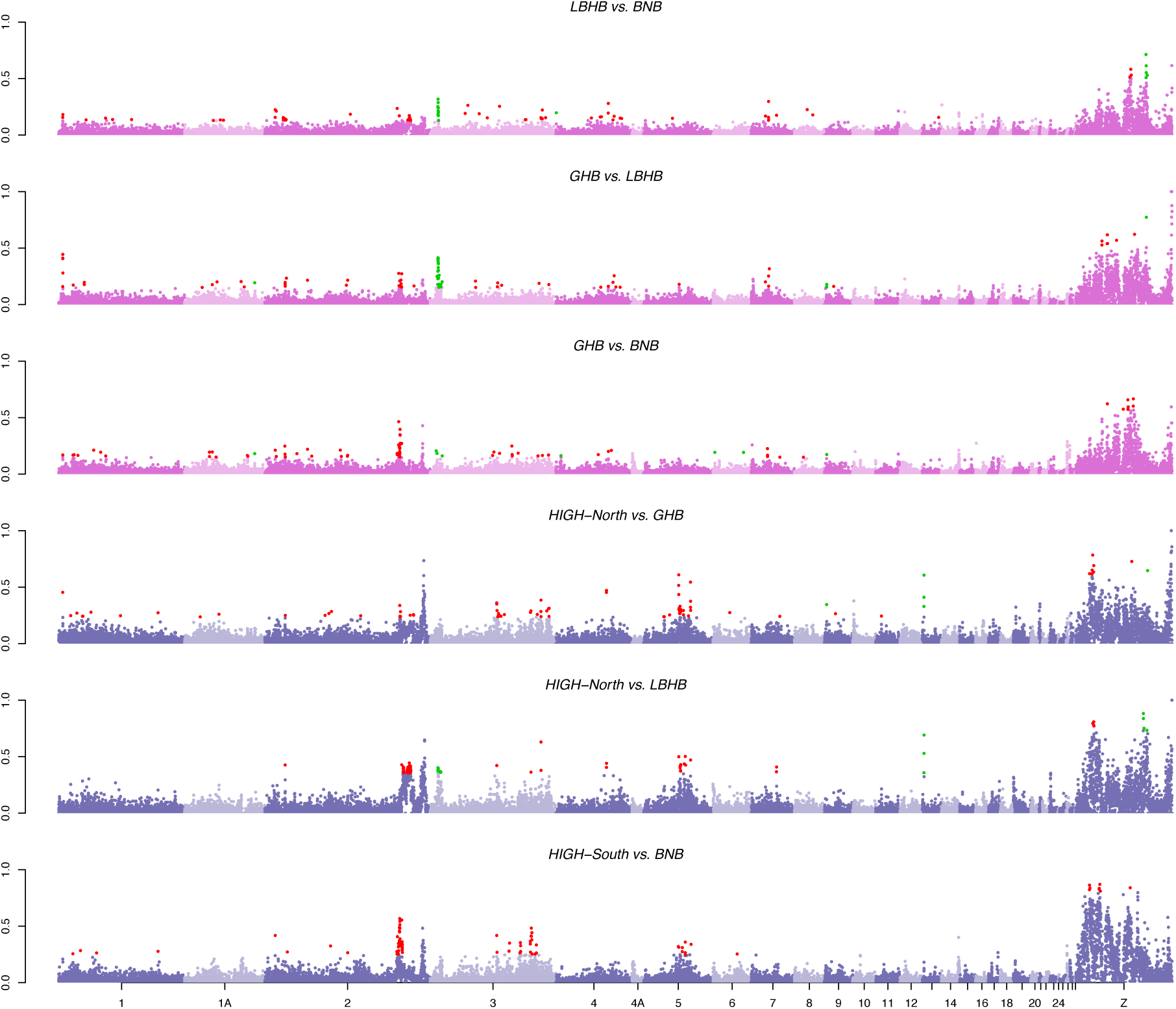
Genomic landscape of *F*_ST_ in the Reunion Grey White-eye. Genomic regions of high *F*_ST_ (top 0.5%) that have a low recombination rate (between 0 and 1cM), for both autosomes and the sex chromosome, are indicated in red. Genomic regions of high *F*_ST_ (top 0.5%) with moderate to high recombination rate (>1cM), that can be viewed as candidate regions for positive selection or reduced gene flow, are indicated in green.

#### 2b Genome-wide screening for selective sweeps

Selective sweeps were detected in both analyses (with or without including a recombination map), but the general values of the CLR were lower when the recombination map was included, and fewer selective sweeps were detected (Figures S5 and S6). The selective sweeps detected in the lowland forms were restricted to the Z chromosome. Specifically, two distinct regions exhibited evidence of selective sweeps in the LBHB form, while one region was identified in both the BNB and GHB forms. Notably, the region containing the selective sweep signal in the BNB form was located near one of the sweep regions identified in the LBHB form. Otherwise, the selective sweeps did not overlap among the different geographical forms. Within the highland form, populations from the two mountain ranges showed marked differences in the location of the sweeps detected: while the HIGH-South form showed two peaks of selective sweeps in the Z chromosome, the selective sweeps identified in the HIGH-North were restricted to a large genomic region at the end of chromosome 2 (Figure 6). Globally selective sweeps do not overlap with regions of elevated differentiation and high recombination except in the LBHB form.

Using gene annotations from the Zebra Finch genome, we identified 261 genes located within candidate sweep regions (Table S5). In lowland forms, candidate sweep regions contained genes associated with immune function and climate-related responses. In highland forms, we detected genes related to altitude adaptation and morphological divergence, including loci potentially linked to bill shape. We also identified a region associated with song behaviour, suggesting possible divergence in vocal traits.

#### 2c Identifying barrier loci

We used a demographic modelling approach for all six pairs of Reunion Grey White-eyes (Figure S7) to infer the intensity of ongoing gene flow between all pairs of forms (model Isolation with Migration) after about 200,000 generations of divergence using sequence loci of 10 kb at most. Effective migration rates were inferred to be homogeneous for all pairs, with the notable exception of a lowland (GHB) vs. a highland form (HIGH-North) where a heterogeneous landscape of gene flow was detected. For this particular pair, we therefore tried to identify barriers to gene flow using this Approximate Bayesian Computation framework. We first defined barrier loci as loci with posterior probability of ‘isolation’ (i.e. no ongoing gene flow) greater than 0.95. Based on this criterion, we identified 28 candidate regions for barrier loci in total, mapping to nine distinct autosomes and the Z chromosome. Genes found in the putative barrier loci were enriched for functions related to altitude adaptation, hypoxia response, immunity, metabolism, pigmentation, and sensory perception, consistent with ecological differentiation and potential reproductive isolation between lowland and highland forms. Genes found in the putative barrier loci were enriched for functions related to altitude adaptation, hypoxia response, immunity, metabolism, pigmentation, and sensory perception, consistent with ecological differentiation and potential reproductive isolation between lowland and highland forms (Table S6).

## Discussion

In this study, we aimed at identifying candidate loci which may have contributed to population divergence and reproductive isolation in a set of comparisons between populations and species along an avian speciation continuum. We used three independent and complementary population genomic approaches: (i) islands of differentiation between populations, taking into account the recombination landscape; (ii) candidate regions for selective sweeps in each geographical form; and (iii) demographic inferences of barriers to gene flow across the genome.

We found that landscapes of differentiation in Mascarene and southern African White-eyes change with the extent of genetic divergence along a speciation continuum. The genomic landscape of differentiation is highly heterogeneous in all comparisons, as already documented in many bird systems (e.g., Poelstra et al. 2014; Irwin et al. 2018; Schield et al. 2025). We observed marked differences in the *F_ST_* distributions in autosomes and the Z chromosome only between recently diverged Reunion Grey White-eye geographical forms. This observation contrasts with the results of a recent study of the silvereye species complex, where pairs of population at the early stage of divergence (150 years only) had similar *F_ST_* distributions in autosomes and Z chromosome (Sendell-Price et al. 2020). This could indicate that some genetic changes on the Z chromosome may be very important to prevent interbreeding in the Reunion Grey White-eye species complex (Bourgeois et al. 2020), and may contribute to reproductive isolation acting as a barrier to gene flow in early stages of divergence. In agreement, introgression has been shown to be reduced on the Z chromosome in several avian hybrid zones, indicating that sex chromosomes may often play an important role in the evolution of reproductive isolation (e.g., Sæther et al. 2007; Carling and Brumfield 2008; Storchová et al. 2010; García et al. 2023). However, as the effective population size of the Z chromosome is ¾ that of autosomes when sex ratio is even (Ellegren 2009), it may also be diverging more rapidly than autosomes solely because of a strong genetic drift (fast-Z effect, e.g., Mank et al. 2007; Fraïsse and Sachdeva 2020), but in that case we would expect a fast-Z effect in the early stage comparisons in silvereyes, which does not appear to be the case (Sendell-Price et al. 2020). Similarly, Leroy et al. (2021a) found little support for a fast-Z effect in the most recent evolution of the Z chromosome.

By taking recombination into account, we intended to identify regions of high differentiation while disentangling the effects of background selection from those of reproductive barriers. Indeed, peaks that are shared between comparisons involving different species correspond to areas of low recombination, as expected assuming a very high conservation of synteny. When performing the sweep detection analysis, we noted that including the recombination map in the analysis similarly reduces the number of detected windows. These regions may correspond to loci that act as partial barriers to gene flow and contribute to reproductive isolation between forms, even in the absence of recent selective sweeps. We also obtained evidence for regions with both high *F_ST_* and high recombination that are likely to host important genes under positive selection as supported by the sweep detection analyses. The impact of recombination detected within the Réunion Grey White-eye is consistent with the correlation in the landscape of *F_ST_* differentiation estimated among more distant Zosterops species (Doren et al. 2017; Dutoit et al. 2017; Vijay et al. 2017). Using the shared *F_ST_* peaks among independent pairs of Zosterops species allows us to account for potential changes in the recombination landscape since their divergence from the zebra finch ancestor. Using a stringent threshold of the top 1% *F_ST_* regions, we found no overlap between our *F_ST_* outliers and the high-*F_ST_* regions detected in the outgroups. The overlap, however, increases substantially when relaxing the threshold to the top 5%. In this case, 28% of our candidate genes fall within the shared high-*F_ST_* regions. This result suggests that the recombination landscape may have evolved slightly since the divergence from the zebra finch, which would not be surprising (Johnston 2024; McAuley et al. 2024). In the future, it will be interesting to estimate the recombination rate across a broader range of species, including here in different Zosterops forms and species in order to ensure that some of the candidate regions do not arise due to a local lineage-specific reduction in recombination rate rather than selection.

Different selection regimes are expected to operate on the different geographical forms of the Reunion Grey White-eye. When comparing highland and lowland forms, we expected to find loci under natural selection, possibly many loci given the polygenic basis of local adaptation along an altitudinal gradient, in contrast to comparisons between lowland forms, where natural selection should be less prominent as the barriers between them are not ecological (Bertrand et al. 2016; Bourgeois et al. 2020). Because phenotypic differences among lowland forms are mostly related to plumage colour, sexual selection could be explaining the maintenance of differences between lowland forms, and could target only a few loci (Delahaie et al. 2017). Plumage coloration has already been identified as a factor involved in reproductive isolation during the early stages of divergence (Poelstra et al. 2014; Turbek et al. 2021). In our model species, a recent study found signatures of selection both on autosomes and Z chromosome for highland vs lowland comparisons, but signatures of selection on the Z chromosome only for lowland comparisons (Bourgeois et al. 2020). Consistently, we also found that candidate outlier regions were mostly present in the sexual chromosome for lowland forms while they involved both autosomes and the sex chromosome in the highland forms. Outlier regions were located in different regions for all the forms, suggesting that selection may be targeting different genomic regions in all forms, including in the lowland forms. Furthermore, sweeps were not overlapping when comparing populations of the highland form from different mountain ranges. This result may appear to be surprising as the two mountains seem to be characterized by relatively homogeneous ecological conditions and similar vegetation types, yet the habitats between the two mountains might differ in many dimensions other than climate and vegetation (Strasberg et al. 2005), especially from a birds’ perspective. For example, the two mountains may differ from one another in biotic factors such as parasites and pathogens, which may be associated with different selective pressures and potentially could also reduce effective gene flow between mountains if coevolved virulence is low in host populations at a local scale but high in more distant populations lacking recent or regular exposure to these parasites (Ricklefs 2010).

Genomic regions of high differentiation contain numerous genes whose functions point to selection on physiological, immune, and behavioural traits. In lowland forms, several candidates such as *MRPL33*, *RBKS*, and *TTC7A* are linked to hypoxia responses (Liu et al. 2020; Cheng et al. 2021; Saravanan et al. 2021), suggesting adaptation to subtle elevational gradients. Other genes associated with energy metabolism (*CAPN8*; Chikina et al. 2016) and immune defense (*ATRNL1*; Arzik et al. 2022) indicate selection for metabolic efficiency and pathogen resistance under differing environmental pressures. Genes related to pigmentation (*BRE*; Twumasi et al. 2024, SLC7A11; Galván et al. 2017) and neurobehavioral traits (*GFRA1*; Colquitt et al. 2021) suggest divergence in coloration and song behaviour within lowland populations. Comparisons between lowland and highland forms revealed additional genes associated with altitude adaptation and immunity. Candidates such as *EPRS*, *JAK2* and *MRPL33* are linked to hypoxia tolerance and metabolic adjustments to elevation (Yoshino et al. 2012; Singh et al. 2018; Saravanan et al. 2021; Xu et al. 2023), while immune-related loci (*ABLIM3*, *CD274*, *FOSL2*) point to selection on pathogen resistance across altitudinal gradients (Renoux et al. 2020; Amat et al. 2021; Sheppard et al. 2022).

Within candidate sweep regions, several genes stand out as potential targets of sexual and natural selection. *KCNS2*, which influences song behaviour in the Zebra Finch (Lovell et al. 2013), together with *KCNV2*, another gene linked to vocal signalling (Lovell et al. 2013), may underlie vocal divergence among Réunion Grey White-eye forms (Freitas 2025). Assortative mating driven by sexual selection, which might be related to differences in song types, has been hypothesized to be an important driver of divergence between lowland forms, as their geographical limits do not match ecological transitions (Delahaie et al. 2017). Interestingly, we identified genes that could be involved in immunity in the LBHB lowland form, including *ANKRA2*, previously associated with pathogen interactions (Casas et al. 2011), as well as *CD24* and *TBK1*, both linked to immune response (Fang et al. 2010; Shao et al. 2023), and *IFNG*, a gene involved in pathogen resistance (Kuttiyarthu Veetil et al. 2024). This may relate to the differences in parasitic pressures that are experienced by the different precipitation regimes found in the western and eastern coasts, which appear to favour distinct assemblages of vectors and parasites (Cornuault et al. 2013). Finally, we evidenced genes involved in pigmentation (*AIM1*; Soejima et al. 2006 and *ASCC3*; Yang et al. 2025) as well as genes linked to reproductive traits and behavioural changes (*LIN28B*; Cousminer et al. 2013 and *TPH2*; Fujita et al. 2023), which may be subject to sexual selection. In addition, potential targets for natural selection were detected in the highland form, including *LRRC2*, previously linked to altitude adaptation in Tibetan pigs (Huang et al. 2020). Finally, *VPS13B*, associated with bill morphology in Darwin finches (Lamichhaney et al. 2015), may underlie the larger beak size of highland individuals (Cornuault et al. 2015). The same gene was also found to be a candidate for genomic differentiation in populations of the silvereye (genus *Zosterops*) (Sendell-Price et al. 2020).

Demographic inferences only detected barriers to gene flow in one lowland vs highland comparison, suggesting that barriers are stronger and/or more numerous between lowland and highland forms, even though the highland form was shown to be of recent origin relative to the history of the complex (Gabrielli et al. 2020). The very low level of differentiation between lowland forms may also limit the power to detect heterogeneity in gene flow between loci. The shift to a drastically different habitat associated with the colonisation of the highlands may have been associated with ecological opportunities that have been increasingly invoked to explain how the availability of contrasting ecological niches or environments can lead to evolutionary diversification (Schluter 2015; Stroud and Losos 2016). These barriers contain genes linked to traits that may directly contribute to reproductive isolation, including altitude adaptation, immunity, and pigmentation, highlighting how natural and sexual selection may limit gene flow between populations. Several genes are associated with altitude and hypoxia tolerance, such as *APTX*, *KDM4C*, *MIPOL1*, *PARD3*, *QARS*, *TRHDE*, and *USP19* (Altun et al. 2012; Salminen et al. 2016; Kang et al. 2017; Edea et al. 2019; Zhang et al. 2019; Wang et al. 2020; Chen et al. 2022), supporting adaptations to reduced oxygen, cooler temperatures, and increased UV radiation in highlands. Genes involved in metabolism and cellular maintenance, i.e. *ABCG1* (cholesterol uptake), *PLGRKT* (adipose function), and *SLC25A20* (lipid usage, possible pigmentation role), may reflect divergent energetic demands across elevations (Storti et al. 2019; Olsen et al. 2021; Miles et al. 2023). Immune-related loci including *CD274*, *DDX58*, and *PDCD1LG2*, suggest differences in pathogen pressures between environments (Sato et al. 2015; Amat et al. 2021; Liu et al. 2023). Visual and sensory adaptation may involve *CNGA3* and potentially *ANKRD32* (UV protection) (Ma et al. 2013; Liu et al. 2018), while *UBE3A* points to divergence in circadian regulation production (Straub et al. 2020). Genes with putative pigmentation effects (*PARD3*, *RIC1*, and *SLC25A20*; Guo et al. 2022; Condori et al. 2025; Zhou et al. 2025) may underlie plumage differences and influence visual signalling. Together, these genes suggest that natural selection on environment-specific physiology and sexual selection related to immunity and pigmentation may be shaping genomic differentiation and potentially strengthening reproductive isolation between lowland and highland forms.

Together, our results demonstrate that integrating recombination variation, selective sweep detection, and demographic inference provides a comprehensive view of how the genomic architecture of divergence evolves and how this can lead to reproductive isolation in the face of gene flow. They also confirm that interpreting patterns of genomic differentiation requires great caution, as already pointed out by other researchers (Duranton et al. 2018), and that explicit consideration of recombination landscapes is an important prerequisite to such analyses. Finally, our study highlights the potential roles of both sexual and natural selection in driving quickly divergence at barrier loci. In line with this result, we identified multiple candidate barrier loci containing genes with varying functions suggesting that mixes of premating and postmating isolating barriers are already at play in this system. This provides further evidence that reproductive isolation can evolve rapidly in some fast-speciating lineages and help understand what underpins the “great speciator paradox” (Diamond et al. 1976).

## Data availability

Raw sequence reads of whole genomes are deposited in the SRA (BioProject PRJNA661201 and PRJEB18566).

## Author contributions

B.M., C.T. and B.N. initiated and coordinated the project. M.G., B.M., C.T. and B.N. conceived the study and designed the experiments. B.M. and C.T. conducted the fieldwork, with the assistance of M.G., T.L. and B.N. M.G. performed the analyses. M.G., T.L., C.R., B.M., C.T. and B.N. interpreted the analyses and wrote the manuscript.

## Funding

This work was supported by Centre National de la Recherche Scientifique (CNRS) through a PEPS grant and the SEE-Life program and the Département de La Réunion. The first author was supported by an MESR (Ministère de l’Enseignement Supérieur et de la Recherche) PhD scholarship during this study.

## Conflict of interest

The authors declare no conflict of interest.

## Acknowledgments

We thank Thomas Duval, Jennifer Devillechabrolle, Guillaume Gélinaud, Marie Manceau, Juli Broggi, Josselin Cornuault, Yann Bourgeois, Joris Bertrand, Boris Delahaie, Dominique Strasberg, Maya Mould, Philipp Heeb, René-Claude Billot, Ben Warren and Jean-Michel Probst for their assistance in the field in Reunion and Mauritius; Reunion National Park and Mauritius National Parks and Conservation Service for granting us permission to conduct fieldwork in protected areas of Reunion and Mauritius, respectively; Benoit Lequette (Reunion National Park) and Vikash Tatayah (Mauritius Wildlife Foundation) for facilitating collecting permits; the provincial authorities in the Free State (South Africa) for granting us permission to collect samples and specimens (permit 01-24158) and Dawie de Swardt (National Museum Bloemfontein) and Jérome Fuchs (MNHN, Paris) for help with organizing field work on African White-eyes. We thank Amaya Iribar and Uxue Suescun for help with the laboratory work; the Genotoul bioinformatics platform for the computational resources they provided and in particular Marie-Stephane Trotard for bioinformatics support.

## Permits

All manipulations on birds were conducted under research permit #602 issued by the Centre de Recherches sur la Biologie des Populations d’Oiseaux (CRBPO), Muséum National d’Histoire Naturelle (Paris).

## Notes

### Competing Interest Statement

The authors have declared no competing interest.

