## Supplementary Material for "Genomics of speciation in a great speciator (Aves: *Zosterops*) reveals the roles of both natural and sexual selection"

**Table S1**. Details about the 33 localities used for genomic analyse on the five White-eye species, including the region, latitude, longitude, and elevation in metres of each sampling locality.

| Sampling site | Region | Latitude | Longitude | Elevation |
|---|---|---|---|---|
| Grande Chaloupe | Reunion | -20,9005 | 55,3758 | 35 |
| Moka | Reunion | -20,9279 | 55,5150 | 219 |
| Rivière du Mât les Bas | Reunion | -20,9794 | 55,6823 | 42 |
| Sentier Ste-Marguerite | Reunion | -21,1088 | 55,6883 | 550 |
| Trois Citernes | Reunion | -21,1589 | 55,7618 | 846 |
| Coulée 2007 | Reunion | -21,2782 | 55,7916 | 194 |
| Basse Vallée | Reunion | -21,3409 | 55,7090 | 687 |
| Piton de l'Entonnoir | Reunion | -21,3566 | 55,6145 | 430 |
| Rivière d'Abord | Reunion | -21,3254 | 55,4949 | 160 |
| L'Etang-salé les Bains | Reunion | -21,2585 | 55,3379 | 68 |
| St-Leu | Reunion | -21,1374 | 55,2953 | 509 |
| Ermitage | Reunion | -21,0733 | 55,2327 | 60 |
| Petit Bernica | Reunion | -21,0254 | 55,2803 | 306 |
| Maïdo | Reunion | -21,0728 | 55,3774 | 2062 |
| Tévelave | Reunion | -21,1689 | 55,3871 | 1962 |
| Bébour | Reunion | -21,0965 | 55,5495 | 1540 |
| Bois Ozoux | Reunion | -21,1979 | 55,6465 | 2283 |
| Pas de Bellecombe | Reunion | -21,2172 | 55,6877 | 2246 |
| Anse des Cascades | Reunion | -21,1837 | 55,8301 | 55 |
| Canot | Reunion | -21,2240 | 55,4131 | 690 |
| Roche Verre Bouteille | Reunion | -20,9877 | 55,3951 | 1251 |
| Grand Matarum | Reunion | -21,1244 | 55,4787 | 1460 |
| Nez de Bœuf | Reunion | -21,2062 | 55,6189 | 2070 |
| Ravine Petit St-Pierre | Reunion | -21,1850 | 55,6722 | 2076 |
| Cap Malheureux | Mauritius | -19,9893 | 57,6284 | 21 |
| Roches Noires forest | Mauritius | -20,1153 | 57,7357 | 15 |
| Le Bouchon | Mauritius | -20,4709 | 57,6786 | 15 |
| Bel Ombre Forest | Mauritius | -20,4735 | 57,4207 | 278 |
| Le Morne Brabant | Mauritius | -20,4598 | 57,3323 | 9 |
| Black River Gorges | Mauritius | -20,3836 | 57,4199 | 117 |
| Macchabé–Brise Fer forest | Mauritius | -20,3787 | 57,4407 | 600 |
| Yemen | Mauritius | -20,3432 | 57,4138 | 164 |
| Le Pouce Mt | Mauritius | -20,1990 | 57,5238 | 605 |
| Sandymount Park | South Africa | -29.75508 | 25.17733 | 1475 |

**Table S2**. Mean $F_{ST}$ in autosomes and the Z chromosome for pairs spanning the continuum of divergence.

| Pair | autosomes | Z chr |
|---|---|---|
| *LBHB vs. BNB* | 0.01 | 0.07 |
| *GHB vs. LBHB* | 0.02 | 0.11 |
| *GHB vs. BNB* | 0.02 | 0.08 |
| *HIGH-North vs. GHB* | 0.03 | 0.14 |
| *HIGH-North vs. LBHB* | 0.03 | 0.16 |
| *HIGH-South vs. BNB* | 0.03 | 0.2 |
| *borbonicus vs. mauritianus* | 0.11 | 0.14 |
| *borbonicus vs. olivaceus* | 0.2 | 0.15 |
| *borbonicus vs. pallidus* | 0.44 | 0.42 |
| *borbonicus vs. virens* | 0.57 | 0.54 |

**Table S3**. Skewness in $F_{ST}$ distribution in autosomes and the Z chromosome for pairs spanning the continuum of divergence.

| pair | autosomes | Z chr |
|---|---|---|
| *LBHB vs. BNB* | 3.4 | 2.34 |
| *GHB vs. LBHB* | 4.36 | 1.93 |
| *GHB vs. BNB* | 3.78 | 2.11 |
| *HIGH-North vs. GHB* | 3.47 | 1.68 |
| *HIGH-North vs. LBHB* | 3.96 | 1.39 |
| *HIGH-South vs. BNB* | 3.92 | 1.07 |
| *borbonicus vs. mauritianus* | 2.32 | 1.75 |
| *borbonicus vs. olivaceus* | 1.6 | 2.29 |
| *borbonicus vs. pallidus* | -0.21 | 0.12 |
| *borbonicus vs. virens* | -0.63 | -0.45 |

**Table S4**. Functions of the genes recovered within 50-kb regions flanking outliers regions of elevated differentiation and moderate to high recombination

*See Excel file*

**Table S5**. Functions of the genes recovered within 50-kb regions flanking candidate selective sweep regions in each of the geographical forms of the Reunion Grey White-eye. Genes in bold are genes that are mentioned in the main text.

*See Excel file*

**Table S6**. Functions of the genes recovered within 50-kb regions flanking candidate barrier regions in each of the geographical forms of the Reunion Grey White-eye.

*See Excel file*

**Figure S1**. Distribution of $F_{ST}$ across the divergence continuum, in autosomes (top pannel) and the Z chromosome (bottom pannel). Black vertical bars indicate the mean value.

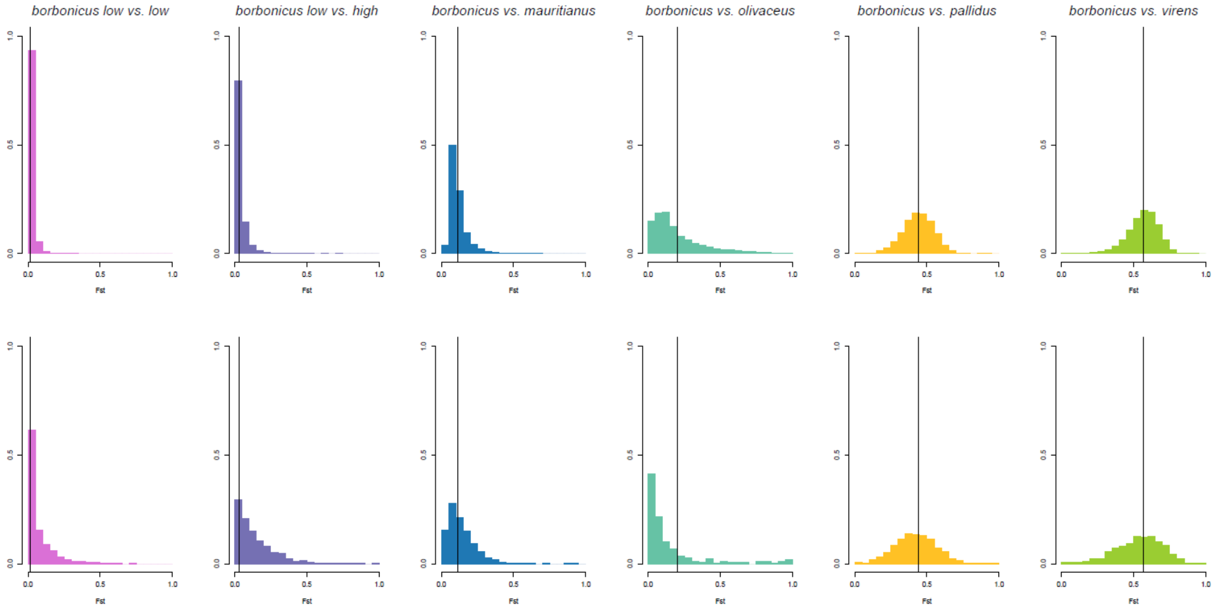

**Figure S2**. Mean $F_{ST}$ per chromosome, with chromosome sorted according to their length, as estimated in the zebra finch genome, from small (left) to large (right) chromosomes. The Z chromosome was removed from these plots.

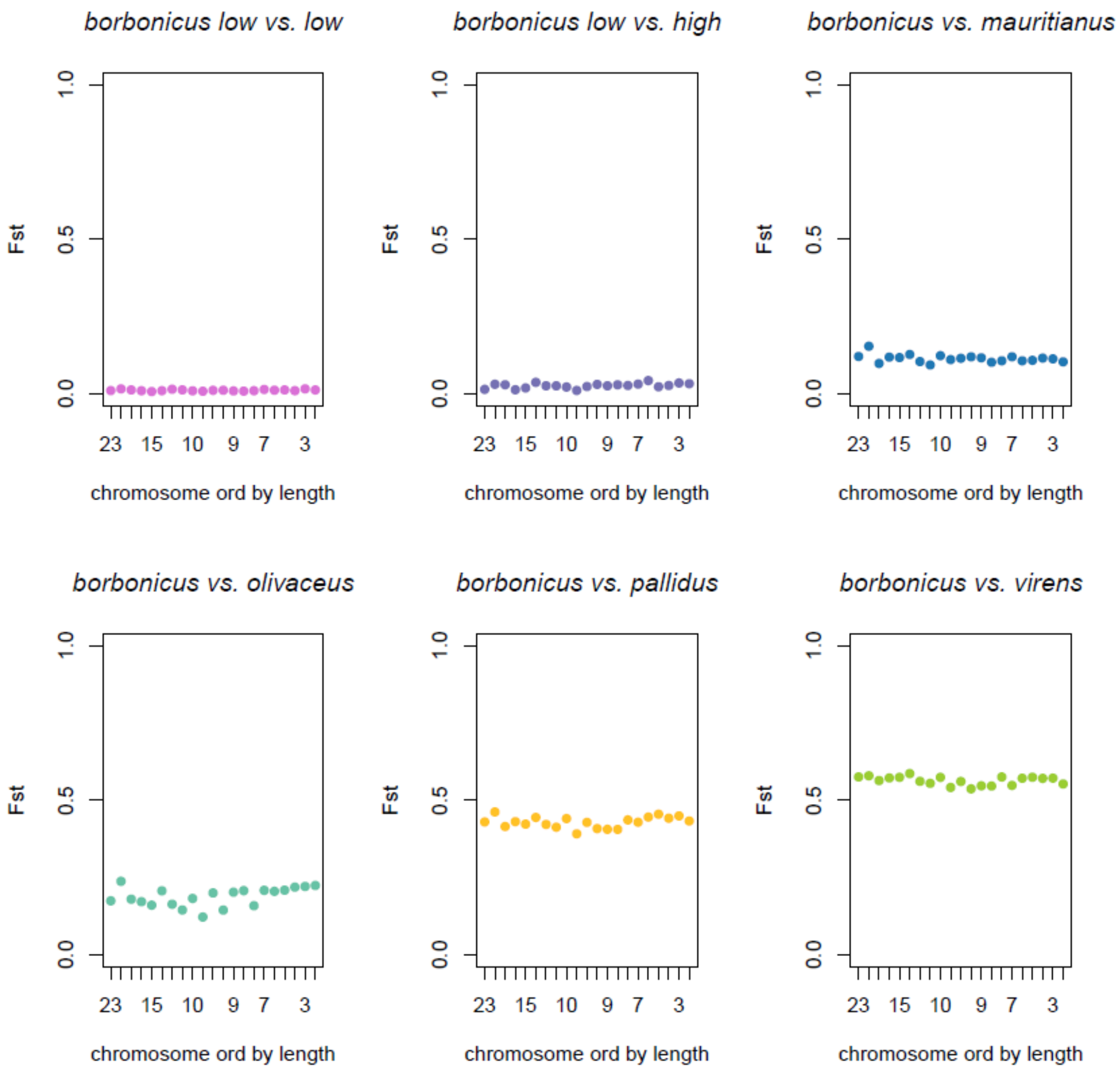

**Figure S3**. Correlations between $F_{ST}$ at the species levels, for autosomes (top pannel) and the Z chromosome (bottom pannel). Top right numbers are pearson correlation coefficients, and the line is a linear regression.

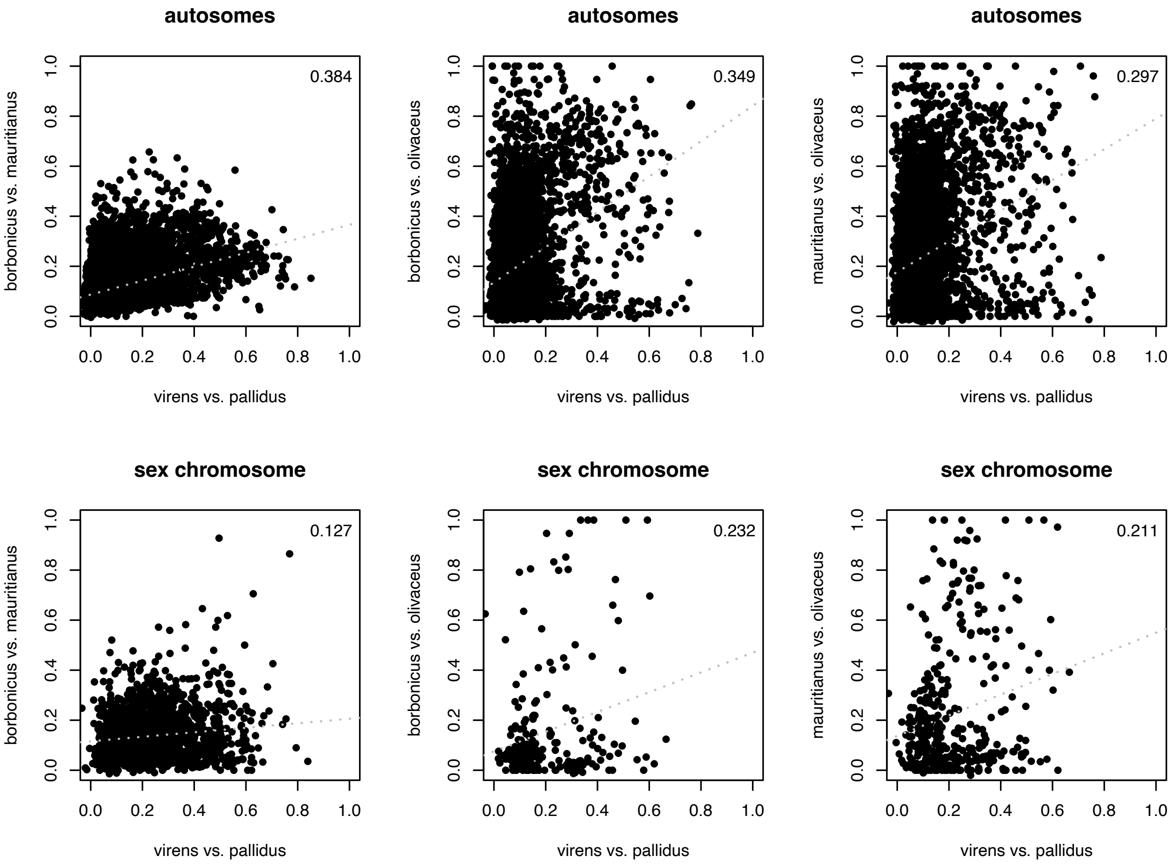

**Figure S4**. $F_{ST}$ according to recombination rate across the divergence continuum, at the species level, in autosomes (top pannel) and the Z chromosome (bottom pannel). Top right numbers are pearson correlation coefficients, and the line is a linear regression.

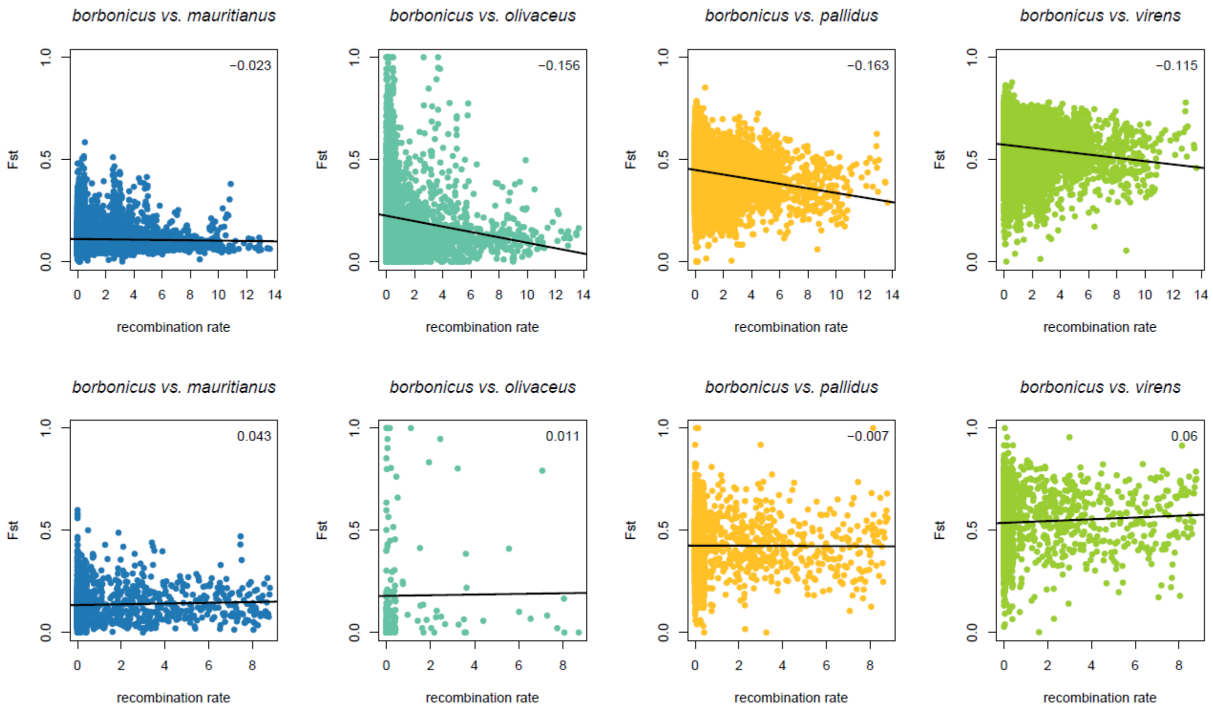

**Figure S5**. Genomic landscape of CLR value from sweepfinder in each Reunion Grey White-eye geographical form, taking into account the recombination landscape. The top 0.1% of the CLR are highlighted in green and were considered as candidate sweeps.

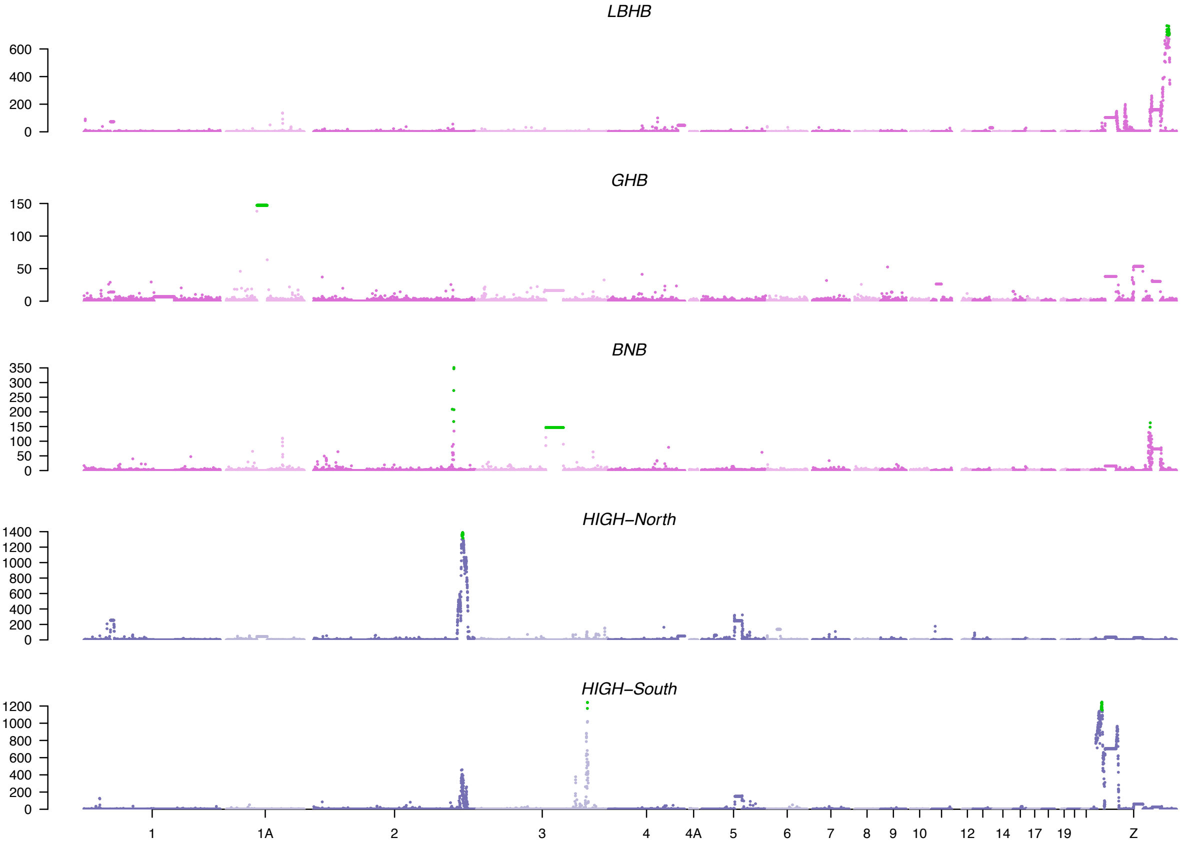

**Figure S6**. Genomic landscape of CLR value from sweepfinder in each Reunion Grey White-eye geographical form, without taking into account the recombination landscape. The top 0.1% of the CLR are highlighted in green.

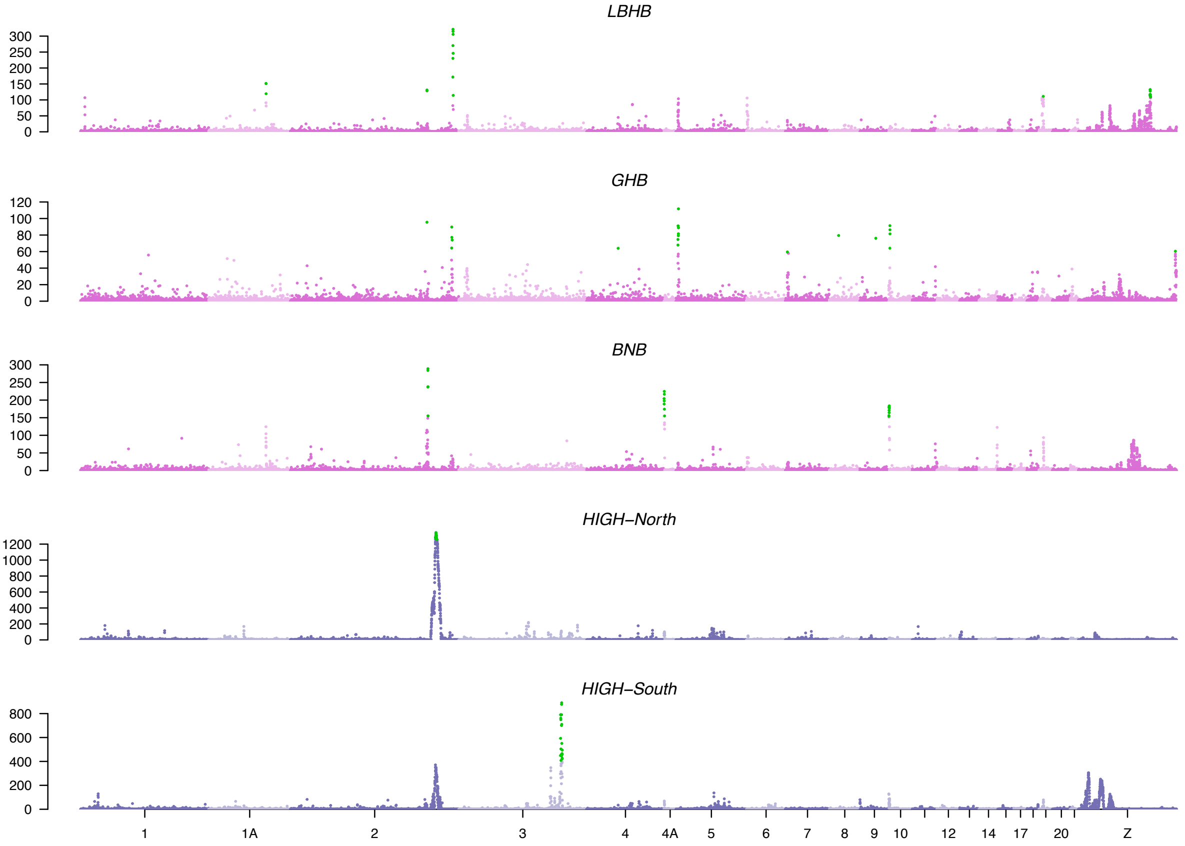

**Figure S7**. Schematic representations of the four divergence models tested for two population models in DILS: Strict Isolation (SI); Isolation with Migration (IM); Ancestral Migration (AM); Secondary Contact (SC).

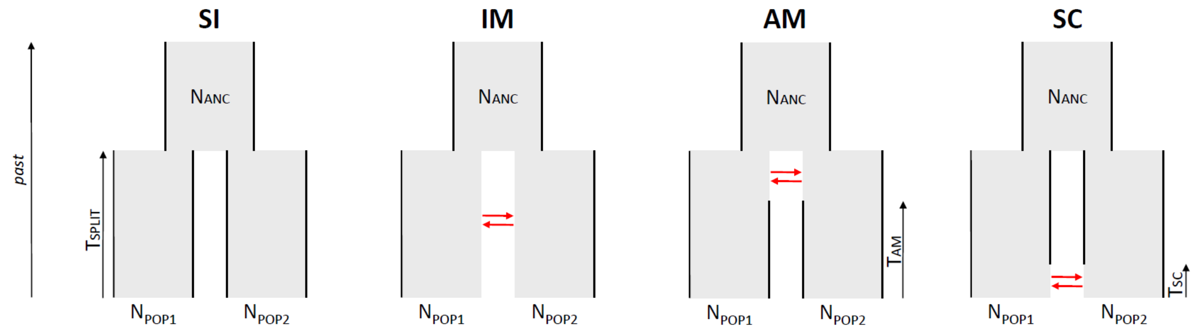

### References supplementary material


Alboali, H., M. H. Moradi, A. H. Khaltabadi Farahani, and H. Mohammadi. 2024. Genome-wide association study for body weight and feed consumption traits in Japanese quail using Bayesian approaches. Poultry Science 103:103208.

Alcaide, M., and S. V. Edwards. 2011. Molecular Evolution of the Toll-Like Receptor Multigene Family in Birds. Mol Biol Evol 28:1703–1715.

Alkorta-Aranburu, G., C. M. Beall, D. B. Witonsky, A. Gebremedhin, J. K. Pritchard, and A. Di Rienzo. 2012. The Genetic Architecture of Adaptations to High Altitude in Ethiopia. PLoS Genet 8:e1003110.

Altun, M., B. Zhao, K. Velasco, H. Liu, G. Hassink, J. Paschke, T. Pereira, and K. Lindsten. 2012. Ubiquitin-specific Protease 19 (USP19) Regulates Hypoxia-inducible Factor 1α (HIF-1α) during Hypoxia*. Journal of Biological Chemistry 287:1962–1969.

Amat, J. A. R., V. Patton, C. Chauché, D. Goldfarb, J. Crispell, Q. Gu, A. M. Coburn, G. Gonzalez, D. Mair, L. Tong, L. Martinez-Sobrido, J. F. Marshall, F. Marchesi, and P. R. Murcia. 2021. Long-term adaptation following influenza A virus host shifts results in increased within-host viral fitness due to higher replication rates, broader dissemination within the respiratory epithelium and reduced tissue damage. PLOS Pathogens 17:e1010174. Public Library of Science.

Anselmi, C. V., A. Malovini, R. Roncarati, V. Novelli, F. Villa, G. Condorelli, R. Bellazzi, and A. A. Puca. 2009. Association of the FOXO3A locus with extreme longevity in a southern Italian centenarian study. Rejuvenation Res 12:95–104.

Arzik, Y., M. Kizilaslan, S. N. White, L. M. W. Piel, and M. U. Çınar. 2022. Genomic Analysis of Gastrointestinal Parasite Resistance in Akkaraman Sheep. Genes 13:2177. Multidisciplinary Digital Publishing Institute.

Bai, D.-P., Y. Chen, Y.-Q. Hu, W.-F. He, Y.-Z. Shi, Q.-M. Fan, R.-T. Luo, and A. Li. 2020. Transcriptome analysis of genes related to gonad differentiation and development in Muscovy ducks. BMC Genomics 21:438.

Bassermann, F., D. Frescas, D. Guardavaccaro, L. Busino, A. Peschiaroli, and M. Pagano. 2008. The Cdc14B-Cdh1-Plk1 Axis Controls the G2 DNA-Damage-Response Checkpoint. Cell 134:256–267. Elsevier.

Benkafadar, N., M. P. Sato, A. H. Ling, A. Janesick, M. Scheibinger, T. A. Jan, and S. Heller. 2024. An essential signaling cascade for avian auditory hair cell regeneration. Developmental Cell 59:280-291.e5. Elsevier.

Bigham, A. W., X. Mao, R. Mei, T. Brutsaert, M. J. Wilson, C. G. Julian, E. J. Parra, J. M. Akey, L. G. Moore, and M. D. Shriver. 2009. Identifying positive selection candidate loci for high-altitude adaptation in Andean populations. Human Genomics 4:79.

Bitaraf Sani, M., M. Mokhtari, Z. Roudbari, O. Karimi, N. Asadzadeh, F. Almathen, and M. H. Banabazi. 2025. The related SNPs and genes to body size using GWAS- latent variable modeling in dromedaries. BMC Genomics 26:645.

Bounas, A., C. Komini, A. Talioura, E.-A. Toli, K. Sotiropoulos, and C. Barboutis. 2023. Adaptive Regulation of Stopover Refueling during Bird Migration: Insights from Whole Blood Transcriptomics. Genome Biol Evol 15:evad061.

Bruders, R., H. V. Hollebeke, E. J. Osborne, Z. Kronenberg, E. Maclary, M. Yandell, and M. D. Shapiro. 2020. A copy number variant is associated with a spectrum of pigmentation patterns in the rock pigeon (*Columba livia*). PLOS Genetics 16:e1008274. Public Library of Science.

Buthasane, W., V. Shotelersuk, W. Chetruengchai, C. Srichomthong, A. Assawapitaksakul, S. Tangphatsornruang, W. Pootakham, C. Sonthirod, S. Tongsima, P. Wangkumhang, A. Wilantho, A. Thongphakdee, S. Sanannu, C. Poksawat, T. Nipanunt, C. Kasorndorkbua, K.-P. Koepfli, B. S. Pukazhenthi, P. Suriyaphol, T. Wongsurawat, P. Jenjaroenpun, and G.

Suriyaphol. 2024. Comprehensive genome assembly reveals genetic diversity and carcass consumption insights in critically endangered Asian king vultures. Sci Rep 14:9455. Nature Publishing Group.
Caizergues, A. E., J. Le Luyer, A. Grégoire, M. Szulkin, J.-C. Senar, A. Charmantier, and C. Perrier. 2022. Epigenetics and the city: Non-parallel DNA methylation modifications across pairs of urban-forest Great tit populations. Evolutionary Applications 15:149–165.
Campana, M. G., A. Corvelo, J. Shelton, T. E. Callicrate, K. L. Bunting, B. Riley-Gillis, F. Wos, J. DeGrazia, E. D. Jarvis, and R. C. Fleischer. 2020. Adaptive Radiation Genomics of Two Ecologically Divergent Hawaiʻian Honeycreepers: The ʻakiapōlāʻau and the Hawaiʻi ʻamakihi. J Hered 111:21–32.
Casas, E., M. D. Garcia, J. E. Wells, and T. P. L. Smith. 2011. Association of single nucleotide polymorphisms in the ANKRA2 and CD180 genes with bovine respiratory disease and presence of Mycobacterium avium subsp. paratuberculosis(1). Anim Genet 42:571–577.
Chen, X., Y. Ji, Y. Cheng, Y. Hao, X. Lei, G. Song, Y. Qu, and F. Lei. 2022. Comparison between short-term stress and long-term adaptive responses reveal common paths to molecular adaptation. iScience 25. Elsevier.
Cheng, J., M. Hui, and Z. Sha. 2019. Transcriptomic analysis reveals insights into deep-sea adaptations of the dominant species, *Shinkaia crosnieri* (Crustacea: Decapoda: Anomura), inhabiting both hydrothermal vents and cold seeps. BMC Genomics 20:388.
Cheng, Y., M. J. Miller, D. Zhang, Y. Xiong, Y. Hao, C. Jia, T. Cai, S.-H. Li, U. S. Johansson, Y. Liu, Y. Chang, G. Song, Y. Qu, and F. Lei. 2021. Parallel genomic responses to historical climate change and high elevation in East Asian songbirds. Proceedings of the National Academy of Sciences 118:e2023918118. Proceedings of the National Academy of Sciences.
Chiba, T., D. Inoue, A. Mizuno, T. Komatsu, S. Fujita, H. Kubota, M. Luisa Tagliaro, S. Park, L. S. Trindade, T. Hayashida, H. Hayashi, H. Yamaza, Y. Higami, and I. Shimokawa. 2009. Identification and characterization of an insulin receptor substrate 4-interacting protein in rat brain: Implications for longevity. Neurobiology of Aging 30:474–482.
Chikina, M., J. D. Robinson, and N. L. Clark. 2016. Hundreds of Genes Experienced Convergent Shifts in Selective Pressure in Marine Mammals. Mol Biol Evol 33:2182–2192.
Coble, D. J., E. E. Sandford, T. Ji, J. Abernathy, D. Fleming, H. Zhou, and S. J. Lamont. 2013. Impacts of Salmonella enteritidis infection on liver transcriptome in broilers. genesis 51:357–364.
Colquitt, B. M., D. P. Merullo, G. Konopka, T. F. Roberts, and M. S. Brainard. 2021. Cellular transcriptomics reveals evolutionary identities of songbird vocal circuits. Science 371:eabd9704. American Association for the Advancement of Science.
Condori, B., M. Recuerda, J. C. Illera, and B. Milá. 2025. The Relative Roles of Selection and Drift in the Chaffinch Radiation (Aves: Fringilla) Across the Atlantic Archipelagos of Macaronesia. Ecology and Evolution 15:e71307.
Cousminer, D. L., D. J. Berry, N. J. Timpson, W. Ang, E. Thiering, E. M. Byrne, H. R. Taal, V. Huikari, J. P. Bradfield, M. Kerkhof, M. M. Groen-Blokhuis, E. Kreiner-Møller, M. Marinelli, C. Holst, J. T. Leinonen, J. R. B. Perry, I. Surakka, O. Pietiläinen, J. Kettunen, V. Anttila, M. Kaakinen, U. Sovio, A. Pouta, S. Das, V. Lagou, C. Power, I. Prokopenko, D. M. Evans, J. P. Kemp, B. St Pourcain, S. Ring, A. Palotie, E. Kajantie, C. Osmond, T. Lehtimäki, J. S. Viikari, M. Kähönen, N. M. Warrington, S. J. Lye, L. J. Palmer, C. M. T. Tiesler, C. Flexeder, G. W. Montgomery, S. E. Medland, A. Hofman, H. Hakonarson, M. Guxens, M. Bartels, V. Salomaa, J. M. Murabito, J. Kaprio, T. I. A. Sørensen, F. Ballester, H. Bisgaard, D. I. Boomsma, G. H. Koppelman, S. F. A. Grant, V. W. V. Jaddoe, N. G. Martin, J. Heinrich, C. E. Pennell, O. T. Raitakari, J. G. Eriksson, G. D. Smith, E. Hyppönen, M.-R. Järvelin, M. I. McCarthy, S. Ripatti, and E. Widén. 2013. Genome-wide association and longitudinal analyses reveal genetic loci linking pubertal height growth, pubertal timing and childhood adiposity. Hum Mol Genet 22:2735–2747.
Cumer, T., A. P. Machado, L. M. San-Jose, A.-L. Ducrest, C. Simon, A. Roulin, and J. Goudet. 2024. The genomic architecture of continuous plumage colour variation in the European barn owl (*Tyto alba*). Proc Biol Sci 291:20231995.

Daneau, G., R. Boidot, P. Martinive, and O. Feron. 2010. Identification of Cyclooxygenase-2 as a Major Actor of the Transcriptomic Adaptation of Endothelial and Tumor Cells to Cyclic Hypoxia: Effect on Angiogenesis and Metastases. Clin Cancer Res 16:410–419.

Davies, S. M. K., M. I. G. Lopez Sanchez, R. Narsai, A.-M. J. Shearwood, M. F. M. Razif, I. D. Small, J. Whelan, O. Rackham, and A. Filipovska. 2012. MRPS27 is a pentatricopeptide repeat domain protein required for the translation of mitochondrially encoded proteins. FEBS Letters 586:3555–3561.

dos Santos, C. G., M. F. Sousa, J. I. G. Vieira, L. R. de Morais, A. A. S. Fernandes, T. de Oliveira Littiere, P. Itajara Otto, M. A. Machado, M. V. G. B. Silva, C. M. Bonafé, A. F. Braga Magalhães, and L. L. Verardo. 2022. Candidate genes for tick resistance in cattle: a systematic review combining post-GWAS analyses with sequencing data. Journal of Applied Animal Research 50:460–470. Taylor & Francis.

Duan, Z., C. Sun, M. Shen, K. Wang, N. Yang, J. Zheng, and G. Xu. 2016. Genetic architecture dissection by genome-wide association analysis reveals avian eggshell ultrastructure traits. Sci Rep 6:28836. Nature Publishing Group.

Dudaniec, R. Y., S. Yadav, J. Catchen, and S. Kleindorfer. 2025. Genomic Introgression Between Critically Endangered and Stable Species of Darwin's Tree Finches on the Galapagos Islands. Evolutionary Applications 18:e70066.

Edea, Z., H. Dadi, T. Dessie, and K.-S. Kim. 2019. Genomic signatures of high-altitude adaptation in Ethiopian sheep populations. Genes Genom 41:973–981.

Eydivandi, S., M. A. Roudbar, M. O. Karimi, and G. Sahana. 2021. Genomic scans for selective sweeps through haplotype homozygosity and allelic fixation in 14 indigenous sheep breeds from Middle East and South Asia. Sci Rep 11:2834. Nature Publishing Group.

Fang, X., P. Zheng, J. Tang, and Y. Liu. 2010. CD24: from A to Z. Cell Mol Immunol 7:100–103. Nature Publishing Group.

Fujita, T., N. Aoki, C. Mori, K. J. Homma, and S. Yamaguchi. 2023. Molecular biology of serotonergic systems in avian brains. Front. Mol. Neurosci. 16. Frontiers.

Galván, I. 2018. Predation risk determines pigmentation phenotype in nuthatches by melanin-related gene expression effects. j. evol. Biol. 31:1760–1771.

Galván, I., Â. Inácio, A. A. Romero-Haro, and C. Alonso-Alvarez. 2017. Adaptive downregulation of pheomelanin-related Slc7a11 gene expression by environmentally induced oxidative stress. Molecular Ecology 26:849–858.

Gao, Y., Y. Lan, H. Liu, and R. Jiang. 2011. The Zinc Finger Transcription Factors Osr1 and Osr2 Control Synovial Joint Formation. Dev Biol 352:83–91.

Goshu, H. A., M. Chu, X. Wu, B. Pengjia, X. Z. Ding, and P. Yan. 2019. Association study and expression analysis of GPC1 gene copy number variation in Chinese Datong yak (*Bos grunniens*) breed. Italian Journal of Animal Science 18:820–832. Taylor & Francis.

Gruber, D. F., J. P. Gaffney, S. Mehr, R. DeSalle, J. S. Sparks, J. Platisa, and V. A. Pieribone. 2015. Adaptive Evolution of Eel Fluorescent Proteins from Fatty Acid Binding Proteins Produces Bright Fluorescence in the Marine Environment. PLOS ONE 10:e0140972. Public Library of Science.

Guo, Q., Y. Jiang, Z. Wang, Y. Bi, G. Chen, H. Bai, and G. Chang. 2022. Genome-Wide Analysis Identifies Candidate Genes Encoding Feather Color in Ducks. Genes 13:1249. Multidisciplinary Digital Publishing Institute.

Gyllenhaal, E. F., S. S. Brady, L. H. DeCicco, A. Naikatini, P. M. Hime, J. D. Manthey, J. Kelly, R. G. Moyle, and M. J. Andersen. 2025. Waves of Colonization and Gene Flow in a Great Speciator. Syst Biol 74:513–525.

Huang, M., B. Yang, H. Chen, H. Zhang, Z. Wu, H. Ai, J. Ren, and L. Huang. 2020. The fine-scale genetic structure and selection signals of Chinese indigenous pigs. Evol Appl 13:458–475.

Jiang, L., D. Bi, H. Ding, X. Wu, R. Zhu, J. Zeng, X. Yang, and X. Kan. 2019. Systematic Identification and Evolution Analysis of Sox Genes in Coturnix japonica Based on Comparative Genomics. Genes 10:314.

Jin, M., H. Wang, G. Liu, J. Lu, Z. Yuan, T. Li, E. Liu, Z. Lu, L. Du, and C. Wei. 2024. Whole-genome resequencing of Chinese indigenous sheep provides insight into the genetic basis underlying climate adaptation. Genet Sel Evol 56:26.

Kang, J., X. Ma, and S. He. 2017. Evidence of high-altitude adaptation in the glyptosternoid fish, Creteuchiloglanis macropterus from the Nujiang River obtained through transcriptome analysis. BMC Evol Biol 17:229.

Kawamura, K., Y. Matsumura, T. Kawamura, H. Araki, N. Hamada, K. Kuramoto, H. Yagi, I. Onoyama, K. Asanoma, and K. Kato. 2024. Endometrial senescence is mediated by interleukin 17 receptor B signaling. Cell Commun Signal 22:363.

Kinoshita, K., K. Tanabe, M. A. Jamal, M. Kyu-Shin, K.-X. Xu, Y.-H. Su, X. Zhang, T. Suzuki, and H.-J. Wei. 2025. Effects of targeted deletion of a 284 bp avian-specific highly conserved element within the Sim1 gene on flight feather development in chickens. Zool Res 46:608–617.

Koivisto, O., A. Hanel, and C. Carlberg. 2020. Key Vitamin D Target Genes with Functions in the Immune System. Nutrients 12:1140. Multidisciplinary Digital Publishing Institute.

Konovalova, S., X. Liu, P. Manjunath, S. Baral, N. Neupane, T. Hilander, Y. Yang, D. Balboa, M. Terzioglu, L. Euro, M. Varjosalo, and H. Tyynismaa. 2018. Redox regulation of GRPEL2 nucleotide exchange factor for mitochondrial HSP70 chaperone. Redox Biology 19:37–45.

Kuttiyarthu Veetil, N., A. E. Henschen, D. M. Hawley, B. Melepat, R. A. Dalloul, V. Beneš, J. S. Adelman, and M. Vinkler. 2024. Varying conjunctival immune response adaptations of house finch populations to a rapidly evolving bacterial pathogen. Front. Immunol. 15. Frontiers.

Laine, V. N., I. Verhagen, A. C. Mateman, A. Pijl, T. D. Williams, P. Gienapp, K. van Oers, and M. E. Visser. 2019. Exploration of tissue-specific gene expression patterns underlying timing of breeding in contrasting temperature environments in a song bird. BMC Genomics 20:693.

Lamichhaney, S., J. Berglund, M. S. Almén, K. Maqbool, M. Grabherr, A. Martinez-Barrio, M. Promerová, C.-J. Rubin, C. Wang, N. Zamani, B. R. Grant, P. R. Grant, M. T. Webster, and L. Andersson. 2015. Evolution of Darwin’s finches and their beaks revealed by genome sequencing. Nature 518:371–375.

Lamichhaney, S., F. Han, J. Berglund, C. Wang, M. S. Almén, M. T. Webster, B. R. Grant, P. R. Grant, and L. Andersson. 2016. A beak size locus in Darwin’s finches facilitated character displacement during a drought. Science 352:470–474.

Lawson, L. P., and K. Petren. 2017. The adaptive genomic landscape of beak morphology in Darwin’s finches. Molecular Ecology 26:4978–4989.

Le Duc, D., G. Renaud, A. Krishnan, M. S. Almén, L. Huynen, S. J. Prohaska, M. Ongyerth, B. D. Bitarello, H. B. Schiöth, M. Hofreiter, P. F. Stadler, K. Prüfer, D. Lambert, J. Kelso, and T. Schöneberg. 2015. Kiwi genome provides insights into evolution of a nocturnal lifestyle. Genome Biology 16:147.

Li, D., Y. Wang, T. Yuan, M. Cao, Y. He, L. Zhang, X. Li, Y. Jiang, K. Li, J. Sun, G. Lv, G. Su, Q. Wang, Y. Pan, X. Li, Y. Jiang, G. Yang, M. A. M. Groenen, M. F. L. Derks, R. Ding, X. Ding, and T. Yu. 2024. Pangenome and genome variation analyses of pigs unveil genomic facets for their adaptation and agronomic characteristics. iMeta 3:e257.

Li, X., A.-M. E. Abdel-Moneim, Z. Hu, N. M. Mesalam, and B. Yang. 2022. Effects of chronic hypoxia on the gene expression profile in the embryonic heart in three Chinese indigenous chicken breeds (Gallus gallus). Front. Vet. Sci. 9. Frontiers.

Liu, C., G. Guo, X. Li, Y. Shen, X. Xu, Y. Chen, H. Li, J. Hao, and K. He. 2023. Identification of novel urine proteomic biomarkers for high stamina in high-altitude adaptation. Front. Physiol. 14. Frontiers.

Liu, X., X. Wang, J. Liu, X. Wang, and H. Bao. 2020. Identifying Candidate Genes for Hypoxia Adaptation of Tibet Chicken Embryos by Selection Signature Analyses and RNA Sequencing. Genes 11:823. Multidisciplinary Digital Publishing Institute.

Liu, Y.-H., L. Wang, T. Xu, X. Guo, Y. Li, T.-T. Yin, H.-C. Yang, Y. Hu, A. C. Adeola, O. J. Sanke, N. O. Otecko, M. Wang, Y. Ma, O. S. Charles, M.-H. S. Sinding, S. Gopalakrishnan, J. Alfredo Samaniego, A. J. Hansen, C. Fernandes, P. Gaubert, J. Budd, P. M. Dawuda, E. Knispel

Rueness, L. Jiang, W. Zhai, M. T. P. Gilbert, M.-S. Peng, X. Qi, G.-D. Wang, and Y.-P. Zhang. 2018. Whole-Genome Sequencing of African Dogs Provides Insights into Adaptations against Tropical Parasites. Mol Biol Evol 35:287–298.

Lovell, P. V., J. B. Carleton, and C. V. Mello. 2013. Genomics analysis of potassium channel genes in songbirds reveals molecular specializations of brain circuits for the maintenance and production of learned vocalizations. BMC Genomics 14:470.

Loyau, T., C. Hennequet-Antier, V. Coustham, C. Berri, M. Leduc, S. Crochet, M. Sannier, M. J. Duclos, S. Mignon-Grasteau, S. Tesseraud, A. Brionne, S. Métayer-Coustard, M. Moroldo, J. Lecardonnel, P. Martin, S. Lagarrigue, S. Yahav, and A. Collin. 2016. Thermal manipulation of the chicken embryo triggers differential gene expression in response to a later heat challenge. BMC Genomics 17:329.

Lugo Ramos, J. S., K. E. Delmore, and M. Liedvogel. 2017. Candidate genes for migration do not distinguish migratory and non-migratory birds. J Comp Physiol A 203:383–397.

Luo, X., J. Guo, J. Zhang, Z. Ma, and H. Li. 2024. Overview of chicken embryo genes related to sex differentiation. PeerJ 12:e17072. PeerJ Inc.

Ma, H., A. Thapa, L. M. Morris, S. Michalakis, M. Biel, M. B. Frank, M. Bebak, and X.-Q. Ding. 2013. Loss of cone cyclic nucleotide-gated channel leads to alterations in light response modulating system and cellular stress response pathways: a gene expression profiling study. Hum Mol Genet 22:3906–3919.

Ma, J., T. Zhang, W. Wang, Y. Chen, W. Cai, B. Zhu, L. Xu, H. Gao, L. Zhang, J. Li, and X. Gao. 2022. Comparative Transcriptome Analyses of Gayal (*Bos frontalis*), Yak (*Bos grunniens*), and Cattle (*Bos taurus*) Reveal the High-Altitude Adaptation. Front. Genet. 12. Frontiers.

Miles, L. A., H. Bai, S. Chakrabarty, N. Baik, Y. Zhang, R. J. Parmer, and F. Samad. 2023. Overexpression of Plg-RKT protects against adipose dysfunction and dysregulation of glucose homeostasis in diet-induced obese mice. Adipocyte 12:2252729. Taylor & Francis.

Mizuno, H., G. Atwal, H. Wang, A. J. Levine, and A. Vazquez. 2010. Fine-scale detection of population-specific linkage disequilibrium using haplotype entropy in the human genome. BMC Genet 11:27.

Mulindwa, J., H. Noyes, H. Ilboudo, L. Pagani, O. Nyangiri, M. P. Kimuda, B. Ahouty, O. F. Asina, E. Ofon, K. Kamoto, J. W. Kabore, M. Koffi, D. M. Ngoyi, G. Simo, J. Chisi, I. Sidibe, J. Enyaru, M. Simuunza, P. Alibu, V. Jamonneau, M. Camara, A. Tait, N. Hall, B. Bucheton, A. MacLeod, C. Hertz-Fowler, E. Matovu, I. Sidibe, D. Mumba, M. Koffi, G. Simo, J. Chisi, V. P. Alibu, A. Macleod, B. Bucheton, C. Hertzfowler, A. Elliot, M. Camara, O. Bishop, J. Mulindwa, O. Nyangiri, M. P. Kimuda, E. Ofon, B. Ahouty, and J. Kabore. 2020. High Levels of Genetic Diversity within Nilo-Saharan Populations: Implications for Human Adaptation. The American Journal of Human Genetics 107:473–486. Elsevier.

Müller, S., A. Perdikari, D. H. Dapito, W. Sun, B. Wollscheid, M. Balaz, and C. Wolfrum. 2020. ESRRG and PERM1 Govern Mitochondrial Conversion in Brite/Beige Adipocyte Formation. Front. Endocrinol. 11. Frontiers.

Mustafi, D., B. M. Kevany, X. Bai, M. Golczak, M. D. Adams, A. Wynshaw-Boris, and K. Palczewski. 2016. Transcriptome analysis reveals rod/cone photoreceptor specific signatures across mammalian retinas. Hum Mol Genet 25:4376–4388.

Obeidat, N. M., M. A. Zihlif, D. A. Alqudah, W. Alshaer, M. Alqaraleh, A. Sharab, and S. S. Abdalla. 2022. Effects of cyclic acute and chronic hypoxia on the expression levels of metabolism related genes in a pancreatic cancer cell line. Biomedical Reports 17:1–11. Spandidos Publications.

Olsen, L., E. Thum, and N. Rohner. 2021. Lipid metabolism in adaptation to extreme nutritional challenges. Developmental Cell 56:1417–1429. Elsevier.

Osipova, E., M.-C. Ko, K. M. Petricek, S. Y. W. Sin, T. Brown, S. Winkler, M. Pippel, J. Jarrells, S. Weiche, M.-B. Mosbech, F. Taborsak-Lines, C. Wang, O. Contreras-Lopez, R.-A. Olsen, P. Ewels, D. Mendez-Aranda, A. Gaede, K. Sadanandan, G. W. Low, A. Monte, N. Ballerstädt, N. M. Adreani, L. Mentesana, A. von Bayern, A. Rico-Guevara, S. Edwards, C. Frankl-Vilches, H.

Kuhl, A. Bakker, M. Gahr, D. Altshuler, W. Buttemer, M. Schupp, M. Baldwin, M. Hiller, and T. Sackton. 2024. Convergent and lineage-specific genomic changes contribute to adaptations in sugar-consuming birds. bioRxiv.

Pease, J. B., R. J. Driver, D. A. de la Cerda, L. B. Day, W. R. Lindsay, B. A. Schlinger, E. R. Schuppe, C. N. Balakrishnan, and M. J. Fuxjager. 2022. Layered evolution of gene expression in “superfast” muscles for courtship. Proceedings of the National Academy of Sciences 119:e2119671119. Proceedings of the National Academy of Sciences.

Qu, Y., C. Chen, Y. Xiong, H. She, Y. E. Zhang, Y. Cheng, S. DuBay, D. Li, P. G. P. Ericson, Y. Hao, H. Wang, H. Zhao, G. Song, H. Zhang, T. Yang, C. Zhang, L. Liang, T. Wu, J. Zhao, Q. Gao, W. Zhai, and F. Lei. 2020. Rapid phenotypic evolution with shallow genomic differentiation during early stages of high elevation adaptation in Eurasian Tree Sparrows. Natl Sci Rev 7:113–127.

Radhakrishnan, S., R. Literman, J. Neuwald, A. Severin, and N. Valenzuela. 2017. Transcriptomic responses to environmental temperature by turtles with temperature-dependent and genotypic sex determination assessed by RNAseq inform the genetic architecture of embryonic gonadal development. PLoS ONE 12:e0172044.

Renoux, F., M. Stellato, C. Haftmann, A. Vogetseder, R. Huang, A. Subramaniam, M. O. Becker, P. Blyszczuk, B. Becher, J. H. W. Distler, G. Kania, O. Boyman, and O. Distler. 2020. The AP1 Transcription Factor Fosl2 Promotes Systemic Autoimmunity and Inflammation by Repressing Treg Development. Cell Reports 31:107826.

Richards, E. J., J. A. McGirr, J. R. Wang, M. E. St. John, J. W. Poelstra, M. J. Solano, D. C. O’Connell, B. J. Turner, and C. H. Martin. 2021. A vertebrate adaptive radiation is assembled from an ancient and disjunct spatiotemporal landscape. Proceedings of the National Academy of Sciences 118:e2011811118. Proceedings of the National Academy of Sciences.

Roberts, R. B., Y. Hu, R. C. Albertson, and T. D. Kocher. 2011. Craniofacial divergence and ongoing adaptation via the hedgehog pathway. PNAS 108:13194–13199. National Academy of Sciences.

Roffler, G. H., S. J. Amish, S. Smith, T. Cosart, M. Kardos, M. K. Schwartz, and G. Luikart. 2016. SNP discovery in candidate adaptive genes using exon capture in a free-ranging alpine ungulate. Molecular Ecology Resources 16:1147–1164.

Romanov, M. N., A. S. Abdelmanova, V. I. Fisinin, E. A. Gladyr, N. A. Volkova, O. A. Koshkina, A. N. Rodionov, A. N. Vetokh, I. V. Gusev, D. V. Anshakov, O. I. Stanishevskaya, A. V. Dotsev, D. K. Griffin, and N. A. Zinovieva. 2023. Selective footprints and genes relevant to cold adaptation and other phenotypic traits are unscrambled in the genomes of divergently selected chicken breeds. J Animal Sci Biotechnol 14:35.

Ruiz-Larrañaga, O., J. Langa, F. Rendo, C. Manzano, M. Iriondo, and A. Estonba. 2018. Genomic selection signatures in sheep from the Western Pyrenees. Genet Sel Evol 50:9.

Salminen, A., K. Kaarniranta, and A. Kauppinen. 2016. Hypoxia-Inducible Histone Lysine Demethylases: Impact on the Aging Process and Age-Related Diseases. Aging Dis 7:180–200.

Saravanan, K. A., M. Panigrahi, H. Kumar, B. Bhushan, T. Dutt, and B. P. Mishra. 2021. Genome-wide analysis of genetic diversity and selection signatures in three Indian sheep breeds. Livestock Science 243:104367.

Sato, H., H. Oshiumi, H. Takaki, H. Hikono, and T. Seya. 2015. Evolution of the DEAD box helicase family in chicken: chickens have no DHX9 ortholog. Microbiology and Immunology 59:633–640.

Sendell-Price, A. T., K. C. Ruegg, E. C. Anderson, C. S. Quilodrán, B. M. V. Doren, V. L. Underwood, T. Coulson, and S. M. Clegg. 2020. The Genomic Landscape of Divergence Across the Speciation Continuum in Island-Colonising Silvereyes (*Zosterops lateralis*). G3: Genes, Genomes, Genetics, doi: 10.1534/g3.120.401352. G3: Genes, Genomes, Genetics.

Shao, Q., F. Fu, P. Zhu, M. Xu, J. Wang, Z. Wang, Y. Yan, H. Wang, J. Ma, Y. Cheng, and J. Sun. 2023. Pigeon TBK1 is involved in antiviral innate immunity by mediating IFN activation. Developmental & Comparative Immunology 147:104758.

Sheppard, E. C., C. A. Martin, C. Armstrong, C. González-Quevedo, J. C. Illera, A. Suh, L. G. Spurgin, and D. S. Richardson. 2022. Genomic associations with poxvirus across divergent island populations in Berthelot's pipit. Molecular Ecology 31:3154–3173.

Singh, D., V. Swarup, H. Le, and V. Kumar. 2018. Transcriptional Signatures in Liver Reveal Metabolic Adaptations to Seasons in Migratory Blackheaded Buntings. Front. Physiol. 9. Frontiers.

Soejima, M., H. Tachida, T. Ishida, A. Sano, and Y. Koda. 2006. Evidence for Recent Positive Selection at the Human AIM1 Locus in a European Population. Mol Biol Evol 23:179–188.

Stonehouse, J. C., L. G. Spurgin, V. N. Laine, M. Bosse, The Great Tit HapMap Consortium, M. A. M. Groenen, K. van Oers, B. C. Sheldon, M. E. Visser, and J. Slate. 2024. The genomics of adaptation to climate in European great tit (*Parus major*) populations. Evol Lett 8:18–28.

Storti, F., K. Klee, V. Todorova, R. Steiner, A. Othman, S. van der Velde-Visser, M. Samardzija, I. Meneau, M. Barben, D. Karademir, V. Pauzuolyte, S. L. Boye, F. Blaser, C. Ullmer, J. L. Dunaief, T. Hornemann, L. Rohrer, A. den Hollander, A. von Eckardstein, J. Fingerle, C. Maugeais, and C. Grimm. 2019. Impaired ABCA1/ABCG1-mediated lipid efflux in the mouse retinal pigment epithelium (RPE) leads to retinal degeneration. eLife 8:e45100. eLife Sciences Publications, Ltd.

Straub, J., A. Gregor, T. Sauerer, A. Fliedner, L. Distel, C. Suchy, A. B. Ekici, F. Ferrazzi, and C. Zweier. 2020. Genetic interaction screen for severe neurodevelopmental disorders reveals a functional link between Ube3a and Mef2 in *Drosophila melanogaster*. Sci Rep 10:1204. Nature Publishing Group.

Twumasi, G., H. Wang, Y. Xi, J. Qi, L. Li, L. Bai, and H. Liu. 2024. Genome-Wide Association Studies Reveal Candidate Genes Associated with Pigmentation Patterns of Single Feathers of Tianfu Nonghua Ducks. Animals 14:85. Multidisciplinary Digital Publishing Institute.

Valero, K. C. W., R. Pathak, I. Prajapati, S. Bankston, A. Thompson, J. Usher, and R. D. Isokpehi. 2014. A candidate multimodal functional genetic network for thermal adaptation. PeerJ 2:e578. PeerJ Inc.

Videvall, E., C. K. Cornwallis, V. Palinauskas, G. Valkiūnas, and O. Hellgren. 2015. The Avian Transcriptome Response to Malaria Infection. Mol Biol Evol 32:1255–1267.

Volkova, N. A., M. N. Romanov, N. Y. German, P. V. Larionova, A. N. Vetokh, L. A. Volkova, A. A. Sermyagin, A. V. Shakhin, D. K. Griffin, J. Sölkner, J. McEwan, R. Brauning, and N. A. Zinovieva. 2025. Genome-Wide Association Studies in Japanese Quails of the F2 Resource Population Elucidate Molecular Markers and Candidate Genes for Body Weight Parameters. International Journal of Molecular Sciences 26:8243. Multidisciplinary Digital Publishing Institute.

Vona, B., R. Maroofian, E. Bellacchio, M. Najafi, K. Thompson, A. Alahmad, L. He, N. Ahangari, A. Rad, S. Shahrokhzadeh, P. Bahena, F. Mittag, F. Traub, J. Movaffagh, N. Amiri, M. Doosti, R. Boostani, E. Shirzadeh, T. Haaf, D. Diodato, M. Schmidts, R. W. Taylor, and E. G. Karimiani. 2018. Expanding the clinical phenotype of IARS2-related mitochondrial disease. BMC Med Genet 19:196.

Walsh, J., G. V. Clucas, M. D. MacManes, W. K. Thomas, and A. I. Kovach. 2019. Divergent selection and drift shape the genomes of two avian sister species spanning a saline–freshwater ecotone. Ecology and Evolution 9:13477–13494.

Wang, C., S.-J. Li, W.-H. Yu, Q.-W. Xin, C. Li, Y.-P. Feng, X.-L. Peng, and Y.-Z. Gong. 2011. Cloning and expression profiling of the VLDLR gene associated with egg performance in duck (*Anas platyrhynchos*). Genet Sel Evol 43:29.

Wang, Q., D. Li, A. Guo, M. Li, L. Li, J. Zhou, S. K. Mishra, G. Li, Y. Duan, and Q. Li. 2020. Whole-genome resequencing of Dulong Chicken reveal signatures of selection. British Poultry Science 61:624–631.

Wang, S., M. J. Ore, E. K. Mikkelsen, J. Lee-Yaw, D. P. L. Toews, S. Rohwer, and D. Irwin. 2021. Signatures of mitonuclear coevolution in a warbler species complex. Nat Commun 12:4279.

Wei, C., H. Wang, G. Liu, F. Zhao, J. W. Kijas, Y. Ma, J. Lu, L. Zhang, J. Cao, M. Wu, G. Wang, R. Liu, Z. Liu, S. Zhang, C. Liu, and L. Du. 2016. Genome-wide analysis reveals adaptation to high altitudes in Tibetan sheep. Sci Rep 6:26770.

Wirthlin, M., N. C. B. Lima, R. L. M. Guedes, A. E. R. Soares, L. G. P. Almeida, N. P. Cavaleiro, G. L. de Morais, A. V. Chaves, J. T. Howard, M. de Melo Teixeira, P. N. Schneider, F. R. Santos, M. C. Schatz, M. S. Felipe, C. Y. Miyaki, A. Aleixo, M. P. C. Schneider, E. D. Jarvis, A. T. R. Vasconcelos, F. Prosdocimi, and C. V. Mello. 2018. Parrot genomes and the evolution of heightened longevity and cognition. Curr Biol 28:4001-4008.e7.

Xu, Y.-X., B. Wang, J.-N. Jing, R. Ma, Y.-H. Luo, X. Li, Z. Yan, Y.-J. Liu, L. Gao, Y.-L. Ren, M.-H. Li, and F.-H. Lv. 2023. Whole-body adipose tissue multi-omic analyses in sheep reveal molecular mechanisms underlying local adaptation to extreme environments. Commun Biol 6:159. Nature Publishing Group.

Yan, Y., H. Zhang, S. Gao, H. Zhang, X. Zhang, W. Chen, W. Lin, and Q. Xie. 2021. Differential DNA Methylation and Gene Expression Between ALV-J-Positive and ALV-J-Negative Chickens. Front. Vet. Sci. 8. Frontiers.

Yang, L., W. Zhao, S. Chen, L. Xue, J. Tian, H. Xu, H. Zhang, H. Wang, Y. Gu, and J. Zhang. 2025. Whole genome resequencing reveals genetic markers for plumage colour in Jingyuan Chicken. Poultry Science 104:105666.

Ye, F., Y. Wang, Q. He, Z. Wang, E. Ma, S. Zhu, H. Yu, H. Yin, X. Zhao, D. Li, H. Xu, H. Li, and Q. Zhu. 2020. Screening of immune biomarkers in different breeds of chickens infected with J subgroup of avian leukemia virus by proteomic. Virulence 11:1158–1176. Taylor & Francis.

Yoshino, S., T. Hara, J. S. Weng, Y. Takahashi, M. Seiki, and T. Sakamoto. 2012. Genetic Screening of New Genes Responsible for Cellular Adaptation to Hypoxia Using a Genome-Wide shRNA Library. PLOS ONE 7:e35590. Public Library of Science.

Zhang, G., C. Cowled, Z. Shi, Z. Huang, K. A. Bishop-Lilly, X. Fang, J. W. Wynne, Z. Xiong, M. L. Baker, W. Zhao, M. Tachedjian, Y. Zhu, P. Zhou, X. Jiang, J. Ng, L. Yang, L. Wu, J. Xiao, Y. Feng, Y. Chen, X. Sun, Y. Zhang, G. A. Marsh, G. Crameri, C. C. Broder, K. G. Frey, L.-F. Wang, and J. Wang. 2013a. Comparative Analysis of Bat Genomes Provides Insight into the Evolution of Flight and Immunity. Science 339:456–460. American Association for the Advancement of Science.

Zhang, L., H. Li, B. Tang, X. Zhao, Y. Wu, T. Jiang, Y. Yao, J. Li, Y. Yao, and L. Wang. 2023. Genomic signatures reveal selection in Chinese and European domesticated geese. Animal Genetics 54:763–771.

Zhang, L., H. Li, X. Zhao, Y. Wu, J. Li, Y. Yao, Y. Yao, and L. Wang. 2024a. Whole genome resequencing reveals the adaptability of native chickens to drought, tropical and frigid environments in Xinjiang. Poultry Science 103:103947.

Zhang, L., J. Liu, F. Zhao, H. Ren, L. Xu, J. Lu, S. Zhang, X. Zhang, C. Wei, G. Lu, Y. Zheng, and L. Du. 2013b. Genome-Wide Association Studies for Growth and Meat Production Traits in Sheep. PLoS ONE 8:e66569.

Zhang, R., R. Li, L. Zhi, Y. Xu, Y. Lin, and L. Chen. 2018. Expression profiles and associations of muscle regulatory factor (MRF) genes with growth traits in Tibetan chickens. British Poultry Science 59:63–67. Taylor & Francis.

Zhang, Y., W. Gou, Y. Zhang, H. Zhang, and C. Wu. 2019. Insights into hypoxic adaptation in Tibetan chicken embryos from comparative proteomics. Comparative Biochemistry and Physiology Part D: Genomics and Proteomics 31:100602.

Zhang, Y., M. Wang, L. Ye, S. Shen, Y. Zhang, X. Qian, T. Zhang, M. Yuan, Z. Ye, J. Cai, X. Meng, S. Qiu, S. Liu, R. Liu, W. Jia, X. Yang, H. Zhang, X. Zhong, and P. Gao. 2024b. HKDC1 promotes tumor immune evasion in hepatocellular carcinoma by coupling cytoskeleton to STAT1 activation and PD-L1 expression. Nat Commun 15:1314. Nature Publishing Group.

Zhong, H.-A., X.-Y. Kong, Y.-W. Zhang, Y.-K. Su, B. Zhang, L. Zhu, H. Chen, X. Gou, and H. Zhang. 2022. Microevolutionary mechanism of high-altitude adaptation in Tibetan chicken populations from an elevation gradient. Evolutionary Applications 15:2100–2112.

Zhou, W., X. Li, X. Zhang, L. Zhu, Y. Peng, C. Zhang, Z. Han, R. Yang, X. Bai, Q. Wang, Y. Zhao, and S. Liu. 2025. Genome-based analysis of the genetic pattern of black sheep in Qira sheep. BMC Genomics 26:114.

Zhou, Z., J. Yang, H. Lv, T. Zhou, J. Zhao, H. Bai, F. Pu, and P. Xu. 2023. The Adaptive Evolution of Leuciscus waleckii in Lake Dali Nur and Convergent Evolution of Cypriniformes Fishes Inhabiting Extremely Alkaline Environments. Genome Biol Evol 15:evad082.

Zhu, H., L. E. Meissner, C. Byrnes, G. Tuymetova, C. J. Tifft, and R. L. Proia. 2020. The Complement Regulator Susd4 Influences Nervous-System Function and Neuronal Morphology in Mice. iScience 23. Elsevier.

Zinzow-Kramer, W. M., B. M. Horton, C. D. McKee, J. M. Michaud, G. K. Tharp, J. W. Thomas, E. M. Tuttle, S. Yi, and D. L. Maney. 2015. Genes located in a chromosomal inversion are correlated with territorial song in white-throated sparrows. Genes, Brain and Behavior 14:641–654.
